# The ICF2 gene Zbtb24 specifically regulates the differentiation of B1 cells *via* promoting heme synthesis

**DOI:** 10.1101/2023.12.23.573176

**Authors:** Jun Wang, He Gao, Sai Zhao, Xiao-Qiu Dai, Xiao-Yuan Qin, Wei-Long Zheng, Can Zhu, Hong-Min Wang, Xue-Mei Zhu, Fang-Yuan Gong, Xiao-Ming Gao, Ying Zhao

**Affiliations:** Institutes of Biology and Medical Sciences, Suzhou Medical College of Soochow University, Suzhou 215123, China; Department of Immunology, School of Biology and Basic Medical Sciences, Suzhou Medical College of Soochow University, Suzhou 215123, China; Department of Pathophysiology, School of Biology and Basic Medical Sciences, Suzhou Medical College of Soochow University, Suzhou 215123, China

**Author notes:** **Corresponding authors:** Dr. Jun Wang and Dr. Xiao-Ming Gao, Institutes of Biology and Medical Sciences, Suzhou Medical College of Soochow University, 199 Ren-Ai Road, Suzhou Industrial Park, Suzhou 215123, Jiangsu Province, China.; or, Dr. Ying Zhao, Department of Pathophysiology, School of Biology and Basic Medical Sciences, Suzhou Medical College of Soochow University, Suzhou 215123, China.

**Keywords:** ICF2, Zbtb24, B1 cells, Plasma cell differentiation, Heme synthesis

## Abstract

Loss-of-function mutations of *ZBTB24* cause the Immunodeficiency, Centromeric Instability and Facial Anomalies syndrome 2 (ICF2). ICF2 is a rare autosomal recessive disorder with immunological defects in serum antibodies and circulating memory B cells, indicating an essential role of ZBTB24 in the terminal differentiation of B cells. Here we generated B-cell specific Zbtb24-deficient mice and systemically investigated its role in B cell development and function both *in vivo* and *in vitro*. Zbtb24 is dispensable for B cell development & maintenance in naive mice. Surprisingly, B-cell specific deletion of Zbtb24 does not evidently compromise germinal center reactions and the resulting primary & secondary antibody responses induced by T-cell dependent antigens, but significantly inhibits T-cell independent antigen-elicited antibody productions *in vivo*. At the cellular level, Zbtb24-deficiency specifically impedes the plasma cell differentiation of B1 cells without impairing their survival, activation and proliferation *in vitro*. Mechanistically, Zbtb24-ablation attenuates heme biosynthesis partially through mTORC1 in B1 cells, and addition of exogenous hemin abrogates the differentiation defects of Zbtb24-null B1 cells. Our study suggests that the defected B1 functions may contribute to recurrent infections in ICF2 patients, and discloses a B1-specific role of Zbtb24 in regulating plasma cell differentiation and antibody production, which is relevant for barrier defenses against invading pathogens.

## INTRODUCTION

Immunodeficiency, centromeric instability and facial anomalies syndrome (ICF [OMIM 242860]) is a rare autosomal recessive and genetically heterogeneous disorder manifested with multiple system failures (Hu et al., 2019; Sterlin et al., 2016). Apart from the hypomethylated pericentromeric regions in specific chromosomes, and varying degrees of facial abnormalities & mental/motor defects, most ICF patients suffer from recurrent respiratory/gastrointestinal infections in early childhood owing to reduced antibody levels (Sterlin et al., 2016; Weemaes et al., 2013). Approximately 50% of ICF cases (ICF1) have mutations in the DNA methyltransferase 3B (*DNMT3B,* OMIM 602900), and about 30% of patients (ICF2) carry nonsense or missense mutations in the zinc finger and BTB domain containing 24 (*ZBTB24,* OMIM 614064) (de Greef et al., 2011; Weemaes et al., 2013; Xu et al., 1999). Although mutations in the cell division cycle associated 7 (*CDCA7*, OMIM 609937), helicase, lymphoid-specific (*HELLS*, OMIM 603946), and ubiquitin-like with plant homeodomain and ring finger domains 1 (*UHRF1*, OMIM 607990) have been recently shown to the associated with ICF3, ICF4 and an atypical ICF case, respectively, there are still few ICF cases (ICFX) left with unidentified disease genes (Thijssen et al., 2015; Unoki et al., 2023). In line with the similar molecular and clinical characteristics across different ICF subtypes, these disease genes act sequentially/corporately to modulate chromatin accessibilities and DNA methylations (Funabiki et al., 2023; Hardikar et al., 2020; Jenness et al., 2018; Ren et al., 2019; Thompson et al., 2018; Unoki et al., 2023; Wu et al., 2016).

Zbtb24 belongs to a large and evolutionary conserved family of transcriptional regulators with about 60 different members, many of which have prominent roles in hematopoietic development, differentiation and function (Cheng et al., 2021; Maeda, 2016; Zhu et al., 2018). For instance, Bcl6 (Zbtb27) promotes germinal center (GC) reactions by driving the differentiation of both GC B cells (GCB) and T follicular helper (Tfh) cells, and thereby is indispensable for the generation of high-affinity memory B cells (Bm) and long-lived antibody-secreting plasma cells (PCs) *in vivo* (Cheng et al., 2021; Toyama et al., 2002; Zhu et al., 2018). In addition, depletion of the ICF4 gene *Hells* in B cell compartment accelerates the premature decay of GCB cells and impairs the generation of high-affinity Bm and antibodies (Cousu et al., 2023). Thus, the absence of GC structures and circulating memory (but not total) B cells in *ZBTB24*-disrupted ICF2 patients indicate that ZBTB24 may possess similar functions in antibody responses *in vivo* (de Greef et al., 2011; Kloeckener-Gruissem et al., 2005; Weemaes et al., 2013; Zhu et al., 2018). Indeed, ZBTB24 modulates the function of human B lymphoblastoid cell line Raji by indirectly repressing Irf4 & Prdm1 in a Bcl6-independent manner (Liang et al., 2016). Moreover, ZBTB24 participates in the nonhomologous end joining process, and thereby promotes the class-switch recombination (CSR) in B cells either directly *via* binding to PARP1 or indirectly by inducing the expression of CDCA7 (Helfricht et al., 2020; Unoki et al., 2019). Nonetheless, the roles of Zbtb24 in the development/function of B cells and *in-vivo* antibody responses remain unknown.

As the effectors of humoral immunity, B cells are comprised of two populations, the predominant B2 subset and the minor B1 compartment (Montecino-Rodriguez and Dorshkind, 2006). B2 cells are continuously produced in the bone marrow (BM) from hematopoietic stem cells (HSCs) throughout life, and the newly formed B cells traffic to the spleen where they eventually mature into either follicular B (FOB) or marginal zone B (MZB) cells. By contrast, B1 cells are mainly seeded during fetal and early neonatal life, and the major population is maintained by self-renewal thereafter, although the adult BM may remain limited potential for B1 development (Baumgarth, 2011; Clarke et al., 2018). Upon activation by T-cell dependent antigens (TD-Ags), the conventional FOB cells enter the GC structure, where they undergo massive proliferation, somatic hypermutation (SHM) & CSR, and ultimately differentiate into Bm or PCs with the capacity to produce high-affinity antibodies (Victora and Nussenzweig, 2022). Conversely, MZB and B1 cells are innate-like B cells with limited B-cell receptor (BCR) repertoire, and are able to rapidly respond to T-cell independent antigens (TID-Ags)/self-antigens by secretion of natural antibodies (Aziz et al., 2015; Baumgarth, 2011; Clarke et al., 2018; Palm and Kleinau, 2021). They promptly and massively differentiate into mature PCs, and thereby provide antibody-mediated innate immune protection during microbial infections (Baumgarth, 2011; Genestier et al., 2007). Given that B1 cells normally reside in the pleura, peritoneum & intestines, they play an important role in barrier immunity to control microbes at mucosal surfaces. Apart from antimicrobial activities, natural antibodies, most of which are produced by B1 cells, help maintain tissue homeostasis by cross-reacting to epitopes expressed on dead/dying autologous cells (Aziz et al., 2015; Baumgarth, 2011; Clarke et al., 2018; Palm and Kleinau, 2021).

It has been well established that the function and fate of B cells are finely controlled by the proper expressions of different transcription factors (TFs). Bcl6/Bach2/Pax5 maintains the phenotype of FOB cells, and promotes their proliferation, SHM and CSR in GC reactions, whereas Prdm1 and Xbp1 augment the differentiation of PCs. Irf4 regulates both of these processes as intermediate Irf4 induces Bcl6 and activation-induced cytidine deaminase (AID), which initiates SHM & CSR, while its high expression upregulates Prdm1 (Ochiai et al., 2013). In addition to the timely & spatially regulated expressions of these TFs, recent studies showed that metabolic reprogramming/adaptation shapes the differentiation & function of B cells as well (Boothby and Rickert, 2017; Foh et al., 2022; Jellusova, 2018). mTORC1 coordinates the increased anabolism and thus plays an essential role in the activation, proliferation and protein synthesis of cells that demand increased energy and building blocks (Boothby et al., 2022). PKC signaling dictates B cell fate by regulating the mitochondrial biogenesis/function partially through mTORC1 signaling (Tsui et al., 2018). Coinciding with their instant actions against pathogens, innate-like B1 and MZB cells exhibit augmented glucose/lipid uptake & metabolism as compared to FOB cells (Jellusova, 2018; Muri et al., 2019). Moreover, high mitochondrial reactive oxygen species (ROS) in activated B cells promotes CSR but dampens the generation of PCs *via* inhibiting the synthesis of heme, which inactivates Bach2 and thus releases its inhibition on Prdm1 expression and PC differentiation (Jang et al., 2015; Watanabe-Matsui et al., 2011).

As the germline deletion of Zbtb24 is embryonic lethal in mice (Wu et al., 2016), we herein generated B-cell specific Zbtb24 knockout mice (Zbtb24^B-CKO^) to investigate its cell-intrinsic role in B cell development and function both *in vitro* and *in vivo*. Our data reveal that cell autonomous Zbtb24 is dispensable for the development/maintenance of B cells in naive mice and the TD-Ag-induced GC reactions and antibody responses, but greatly promotes the PC differentiation of peritoneal B1 cells *via* augmenting mTOCR1 activity and heme biosynthesis upon activation. Thus, our study not only suggests that the barrier defenses against pathogens might be impaired, owing to hampered differentiation of B1 cells, in respiratory & gastrointestinal tracts of ICF2 patients, but also provides mechanistic insights into the specific role of Zbtb24 in regulating B1 cell function.

## RESULTS

### Little effect of Zbtb24-deficiency on the development and phenotype of B cells in mice

Given that the germline knockout of Zbtb24 is embryonic lethal in mice (Wu et al., 2016), we generated mice harboring a conditional *Zbtb24^loxp/+^*allele with two loxp sites flanking the exon 2 which contains the translation starting site (**Figure 1A**). The N-terminal 316 amino acids, including the BTB domain, the AT-hook domain and the first C_2_H_2_ Zinc finger motif, of Zbtb24 protein would be specifically removed in Cre-expressing cells & tissues, which results in a frame shift in the protein coding sequence and thereby generates a premature stop codon. To address cell-intrinsic roles of Zbtb24 in the development and function of B cells *in vivo*, we crossed the *Zbtb24^loxp/+^* mice with the *Cd19*-driven Cre-expressing mouse strain (*Cd19^Cre/+^*) to generate B-cell specific Zbtb24 deficient (Zbtb24^B-CKO^, *Cd19^Cre/+^Zbtb24^loxp/loxp^*) and the littermate control (Cd19^Cre/+^, *Cd19^Cre/+^Zbtb24^+/+^*) mice. Zbtb24^B-CKO^ mice grew normally, and were phenotypically indistinguishable from the control Cd19^Cre/+^ mice.

**Figure 1.**
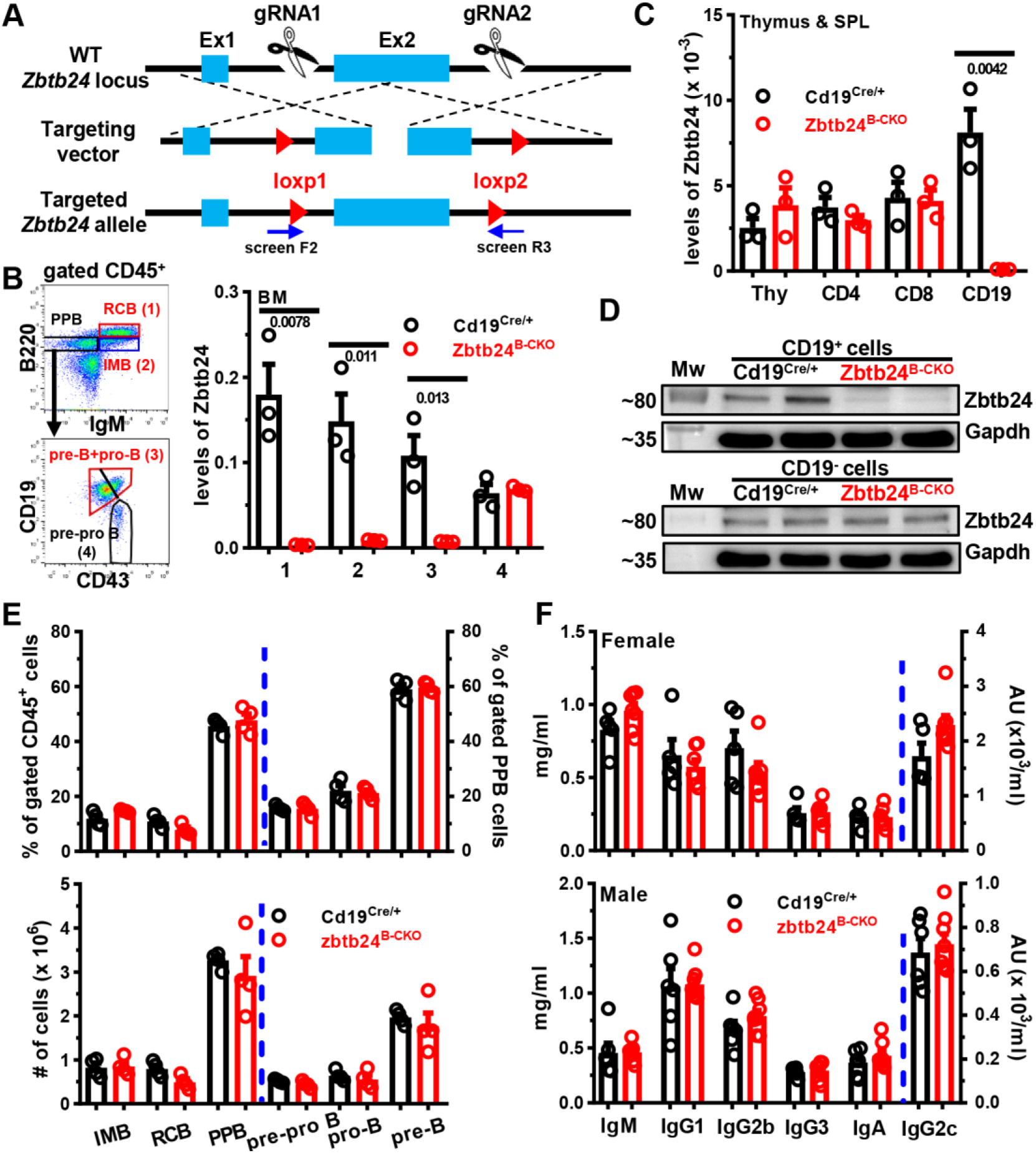
No effect of B-cell specific depletion of Zbtb24 on B-cell development in the bone marrow (BM) and baseline serum antibody levels in mice. **A**, a schematic representation for generating *Zbtb24^loxp/+^* mice using the CRISPER/CAS9 strategy. Screen F2 and R3 denote primers used to select correctly-targeted ES clones. **B-D**, efficient and specific deletions of Zbtb24 in CD19^+^ B cells in Cd19^Cre/+^Zbtb24^loxp/loxp^ (Zbtb24^B-CKO^) mice. Levels of Zbtb24 were analyzed in FACS-sorted B220^++^IgM^+^ recirculating mature B cells (RCB, gate 1), B220^+^IgM^+^ immature B cells (IMB, gate 2), B220^+^IgM^-^CD19^+^CD43^-^ pre-B plus B220^+^IgM^-^CD19^+^CD43^+^ pro-B cells (gate 3), and B220^+^IgM^-^CD19^-^CD43^+^ pre-pro B cells (gate 4) in the BM (**B**), or FACS-purified CD4^+^ (CD4), CD8^+^ (CD8) T cells and CD19^+^ (CD19) B cells in spleens (SPL) as well as total thymocytes (Thy) (**C**) of control Cd19^Cre/+^ *vs.* Zbtb24^B-CKO^ mice by quantitative PCR. Sorting strategies for indicated BM B-cell subsets were shown in the left panel of **B**. Expressions of Zbtb24 were normalized to internal Gapdh (**B&C**). Protein levels of Zbtb24 were analyzed in beads-purified splenic CD19^+^ B cells (*upper panel*) and the remaining CD19^-^ fraction (*lower panel*) by Western blot (**D**). **E**, bar graphs showing the percents (*upper panel*) or absolute numbers (*lower panel*) of indicated B-cell subsets in the BM of control Cd19^Cre/+^ and Zbtb24^B-CKO^ mice. **F**, bar graphs showing levels of total IgM, IgG1, IgG2b, IgG2c, IgG3 and IgA in sera of female (8-10 weeks old) and male (9-11 weeks old) Cd19^Cre/+^ *vs.* Zbtb24^B-CKO^ mice. Each symbol/lane represents a single mouse of the indicated genotype, and numbers below horizonal lines indicate *P* values determined by student *t*-test (**B&C**). PPB, pre-pro/pro/pre B cells; Mw: molecular weight marker; AU, arbitrary units.

In Zbtb24^B-CKO^ mice, Zbtb24 was specifically and efficiently deleted from CD19-expressing pre-B & pro-B cells onwards in the BM and peripheral B cells, while its expression in CD19-negative BM pre-pro B cells, splenocytes and thymocytes were intact as analyzed by Q-PCR and/or western blot (**Figure 1B-D**). Notably, lack of Zbtb24 affected neither the percentages/absolute numbers of different developing stages of B cells in the BM (**Figure E**), nor the numbers of splenic FOB/MZB cells as well as the peritoneal B2, B1a/B1b cells in the periphery (**Figure S1A&B**). Moreover, the percentages of GCB in the spleens, mesenteric lymph nodes (MLN) & Peyer’s Patches (PP), and the serum total IgM, IgG & IgA levels were all normal in Zbtb24^B-CKO^ mice (**Figure S1C**, and **Figure 1F**). Thus, depletion of B-cell Zbtb24 has little impact on the early BM B-cell development, the phenotype & maintenance of peripheral B cells, and serum antibody levels in naive mice.

### Largely normal humoral responses to TD-Ags in Zbtb24^B-CKO^ mice

The major immunological features of ICF2 patients are hypogammaglobulinemia and lack of circulating Bm cells, possibly owing to the disrupted germinal center reactions (Banday et al., 2020; Kloeckener-Gruissem et al., 2005; Liang et al., 2016; Weemaes et al., 2013). We thus first characterized TD-Ag-induced humoral responses in mice with B-cell specific Zbtb24 deficiency. One week after immunization with NP_19_-OVA (4-Hydroxy-3-nitrophenylacetyl hapten conjugated to ovalbumin) emulsified in IFA (incomplete Freund adjuvant, NP-OVA/IFA), serum levels of NP-specific IgM, IgG1 and IgG2c were significantly reduced in Zbtb24^B-CKO^ mice. Surprisingly, on day 14 only NP-specific IgG2c was still slightly decreased, while the other two subtypes were reverted to normal levels in Zbtb24^B-CKO^ mice (**Figure 2A&B**). Moreover, the percentages & numbers of CD19^+^CD95^high^CD38^low^ GCB as well as CD19^low^CD138^++^ PC in spleens were all comparable between control and Zbtb24^B-CKO^ mice (**Figure 2C&D**). Within gated GCB cells, percents of CD86^low^CXCR4^high^ dark zone centroblasts and CD86^high^CXCR4^low^ light zone centrocytes were comparable between the two genotypes of mice as well (**Figure 2C**). Consistent with our previous findings in human cells, lack of Zbtb24 did not affect the expressions of Bcl6, the key driving TF for GC reactions, in gated either GCB (defined as CD19^+^CD95^high^CD38^low^) compartment or total CD19^+^ B cells (**Figure S2A&B**). In addition, the numbers of splenic GCB & PC cells were comparable in Cd19^Cre/+^ & Zbtb24^B-CKO^ mice immunized with sheep red blood cells (SRBC) as well (**Figure S2C**), further corroborating the dispensability of Zbtb24 on their differentiations *in vivo*. Notably, Zbtb24 was efficiently depleted in purified GCB cells (**Figure S2D**), thus it is unlikely that Zbtb24-expressing cells outcompete Zbtb24-null ones to fill the GCB compartment in TD-Ag-immunized Zbtb24^B-CKO^ mice.

**Figure 2.**
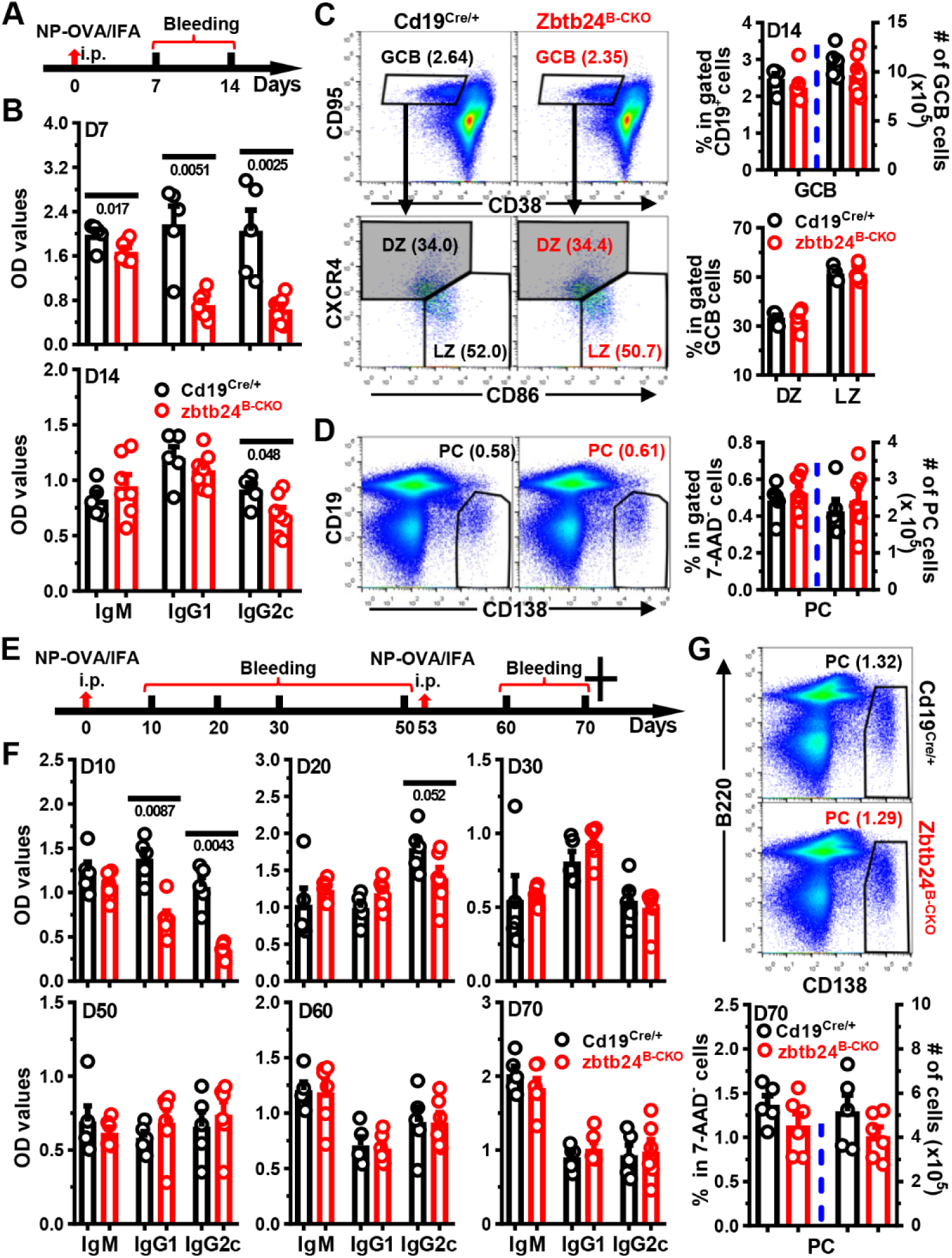
No significant defects in antibody responses in Zbtb24^B-CKO^ mice post primary/secondary NP-OVA/IFA immunization. Mice (10-week old males in **A-D**; and 8-week old females in **E-G**) were intraperitoneally (i.p.) immunized with NP_19_-OVA emulsified in IFA (1:1, 50 μg/100 μl/mouse) on D0, followed by i.p. rechallenging with NP_19_-OVA/IFA (3:1, 20 μg/100 μl/mouse) on D53 (only in **E-G**). Sera were collected on indicated days and NP-specific antibodies were detected by ELISA. Cells in spleens on D14 (**C&D**) or D70 (**G**) were stained with antibodies, in combination with 7-AAD to exclude dead cells, to visualize the percents of CD19^+^CD38^low^CD95^high^ germinal center B (GCB) cells or CD19^-/low^CD138^high^ plasma cells (PC) by flow cytometry. **A&E**, schematic diagrams illustrating the immunization and bleeding schedules of mice. **B&F**, bar graphs showing the optical density (OD) values of NP-specific antibodies in sera of Cd19^Cre/+^ and Zbtb24^B-CKO^ mice on indicated days. **C,** representative pseudo-plots showing the percentages of GCB (within gated 7-AAD^-^CD19^+^ B cells) and frequencies of dark zone centroblasts (DZ, CXCR4^high^CD86^low^) and light zone centrocytes (LZ, CXCR4^low^CD86^high^) within gated GCB cells in control Cd19^Cre/+^ and Zbtb24^B-CKO^ mice. **D&G**, representative pseudo-plots showing the percentages of PCs in spleens of mice on D14 (**D**) or D70 (**G**). Cumulative data on the percentages or absolute numbers of indicated B-cell subsets were shown in the right of **C&D** or bottom of **G**. Each symbol represents a single mouse of the indicated genotype, and numbers below horizonal lines indicate *P* values while those in brackets denote percentages.

Given that the generation of specific antibodies appeared to be delayed in Zbtb24^B-CKO^ mice upon immunization, we immunized mice with NP-OVA/IFA on day 0, and boosted the mice with the same antigen intraperitoneally once again on day 53 to further determine the impact of B-cell Zbtb24-deficiency on humoral responses elicited by TD-Ags *in vivo* (**Figure 2E**). Akin to previous results, control mice produced significantly more NP-specific IgG1 and IgG2c on day 10, and the differences went down greatly on day 20 post the primary immunization (**Figure 2F**). Of note, comparable levels of NP-specific antibodies were detected between the two groups thereafter, even on day 7 & 17 post the secondary challenge (**Figure 2F**). The numbers of PCs were normal in spleens of Zbtb24^B-CKO^ as well (**Figure 2G)**.

We also immunized mice with a high dose of NP_19_-OVA (100 μg/mouse) adsorbed on alum, which predisposes to the Th2 immune response instead of Th1 triggered by IFA *in vivo*. Again, reduced antigen-specific IgG1 were only observed early (day 7) post the primary immunization (**Figure S3A&B**). Moreover, specific deletion of Zbtb24 in B-cell compartment did not affect the ratios of high-affinity to low-affinity (against NP_2_-BSA & NP_25_-BSA, respectively) IgG1 & IgG2c before (i.e., day 70) and two weeks (i.e., day 84) post the secondary immunization (**Figure S3C&D**). Given that Zbtb proteins may act redundantly to maintain the integrity and function of immune cells (Zhu et al., 2018), we conducted RNA-Seq analysis on purified splenic B cells before & after LPS stimulation to determine if any Zbtb proteins were evidently upregulated to compensate for the loss of Zbtb24. Levels of all Zbtb proteins, except Zbtb24, did not differ significantly between time-matched control and Zbtb24^B-CKO^ splenic B cells (**Table S1**).

To exclude any potential off-target deletions of Zbtb24 in Zbtb24^B-CKO^ mice, we next adoptively transferred purified splenic control or Zbtb24-deficient CD19^+^ B cells, together with WT CD4^+^ Th cells, into Rag2^-/-^ recipients (**Figure S4A**). After immunization, serum levels of antigen-specific IgG, and total numbers of splenic CD19^+^ B cells or CD19^low^CD138^++^ PCs did not differ in Rag2^-/-^ recipients implanted with control *vs.* Zbtb24-dificient B cells (**Figure S4C-D**). Notably, the total serum IgM and IgG were reduced in Rag2^-/-^ mice reconstituted with Zbtb24-depleted B cells (**Figure S4B**). Decreased total polyreactive IgM might underlie the reduced levels of NP-specific IgM in Zbtb24^B-CKO^ mice (**Figure S4B&C**), as ablation of Zbtb24 did not profoundly affect the CSR ability of murine splenic B cells *in vitro* (**Figure S5**).

Collectively, these data indicate that B-cell intrinsic Zbtb24 does not play an essential role in the generation and maintenance of GCB cells, their affinity maturation as well as later differentiation toward Bm or PCs after immunization, albeit that Zbtb24^B-CKO^ mice show delayed/mitigated early extrafollicular antibody productions induced by TD-Ag.

### Significantly reduced humoral responses to TID-Ags in Zbtb24^B-CKO^ mice

The reduced total serum IgM & IgG, most of which likely react against autoantigens and commensal gut bacteria, in Rag2^-/-^ mice reconstituted with Zbtb24-deficient B cells indicate that lack of Zbtb24 hampers antibody responses elicited by TID-Ags (**Figure S4B**). Indeed, Zbtb24^B-CKO^ mice produced significantly decreased amounts of NP-specific IgM, IgG1 and IgG3, three major antibody subtypes against soluble protein or carbohydrate antigens in mice (Zhao et al., 2022), in the first two weeks after immunization with the type II TID-Ag NP-Ficoll. On day 21 & 35, levels of IgG1 were still reduced, and the other two antibody subtypes tended to decrease as well (**Figure 3A**). Likewise, significantly less NP-specific IgM, IgG1, IgG3 and IgG2b were detected in male Zbtb24^B-CKO^ mice after immunization with the type I TID-Ag NP-LPS (**Figure 3F**). As expected, reduced percentages/numbers of CD19^low^CD138^++^ PCs were detected in spleens and/or peritoneal cavities (PeC) of Zbtb24^B-CKO^ mice (**Figure 3B-E & G-H**).

**Figure 3.**
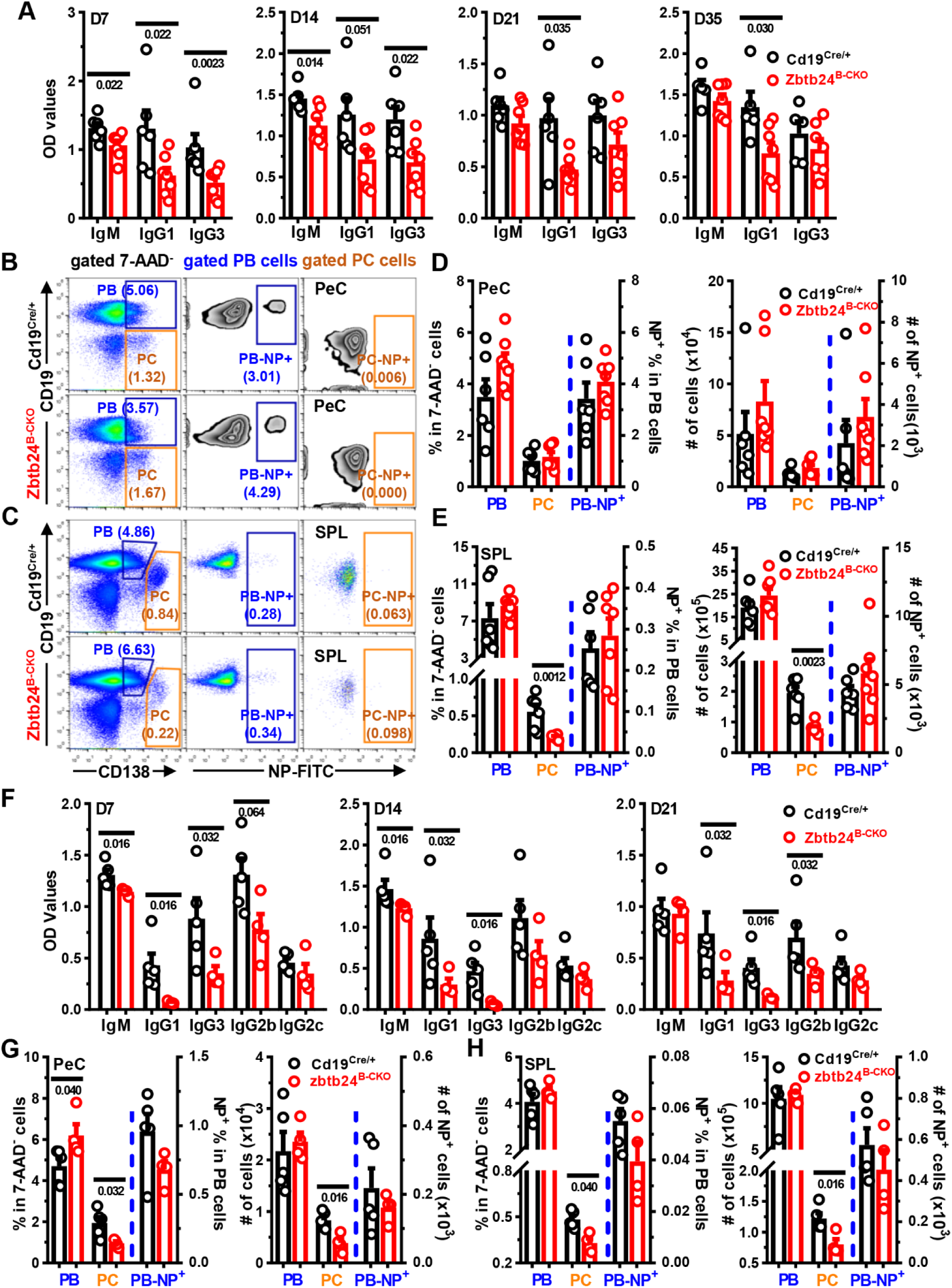
Significantly reduced antigen-specific antibody levels in Zbtb24^B-CKO^ mice after TID-Ag immunization. Mice were intraperitoneally immunized with the type II TID-Ag NP-Ficoll (10 μg/200 μl PBS/mouse, **A-E**) or type I TID-Ag NP-LPS (20 μg/200 μl PBS/mouse, **F-H**) on D0. Blood was taken at indicated times and NP-specific antibodies in sera were determined by ELISA in plates coated with NP_25_-BSA. **A&F**, bar graphs showing the optical density (OD) values of NP-specific antibodies in diluted sera of Cd19^Cre/+^ *vs.* Zbtb24^B-CKO^ mice on indicated days post immunization with NP-Ficoll (**A**) or NP-LPS (**F**). **B&C**, representative pseudo/zebra-plots illustrating the gating strategies for CD19^+^CD138^+^ plasma blasts (PB)/CD19^low^CD138^high^ plasma cells (PC) and percents of NP^+^ cells within gated PB (PB-NP^+^) & PC (PC-NP^+^) cells in peritoneal cavities (PeC, **B**) or spleens (SPL, **C**) of mice on D35 post NP-Ficoll immunization. **D&E and G&H**, bar graphs showing the percentages & absolute numbers of PB & PC cells as well as the NP^+^ cells within gated PB/PC cells (PB-NP^+^/PC-NP^+^, respectively) in PeC (**D&G**) and SPL (**E&H**) of mice on D35 post NP-Ficoll (**D&E**) or on D21 post NP-LPS (**G&H**) immunization. Each symbol represents a single mouse of the indicated genotype (female, 10 weeks of age in **A-E**; and male, 9 weeks of age in **F-H**), and numbers below horizonal lines in bar graphs indicate *P* values.

B1 and MZB cells are considered the main responders to TID-Ags (Haas, 2011). We thus tried to delineate which cell type was responsible for the diminished antibody levels in TID-Ag-immunized Zbt24^B-CKO^ mice *via* detailed flow cytometry analyses. Five weeks post NP-Ficoll immunization, the numbers of NP-specific cells in differentially gated B-cell compartments were all comparable in the spleens and PeC between Cd19^Cre/+^ and Zbtb24^B-CKO^ mice (**Figure S6**), possibly due to the late analyzing time point. Of note, significantly less NP^+^ B1, but not FOB or MZB, cells were observed in the spleens of Zbtb24^B-CKO^ mice three weeks post NP-LPS inoculation, and NP^+^ B1 cells tended to decrease in their PeC as well (**Figure S7**), indicating that B1 was the target B-cell subset through which Zbtb24 promoted TID-Ag-induced antibody responses *in vivo*.

### Zbtb24-deficiency impairs the differentiation and antibody-producing ability of B1 cells *in vivo*

To unambiguously reveal the roles of Zbtb24 in B1 cells, we adoptively transferred peritoneal B1 cells, purified from Cd19^Cre/+^ or Zbtb24^B-CKO^ mice by FACS-sorting, into Rag2^-/-^ mice with no mature T/B cells and serum antibodies (**Figure 4A**). Total IgM levels were significantly lower in Rag2^-/-^ recipients received Zbtb24-deficient B1 cells (**Figure 4B**). Moreover, 7 days after NP-LPS immunization (i.e., day 27 post transfer), Zbtb24-deficient B1-reconstituted Rag2^-/-^ recipients produced significantly less NP-specific IgM than those harboring control Cd19^Cre/+^ B1 cells (**Figure 4C**). Total or NP-specific IgG was hardly detectable in recipients implanted with Zbtb24-deficient or sufficient B1 cells (data not shown).

**Figure 4.**
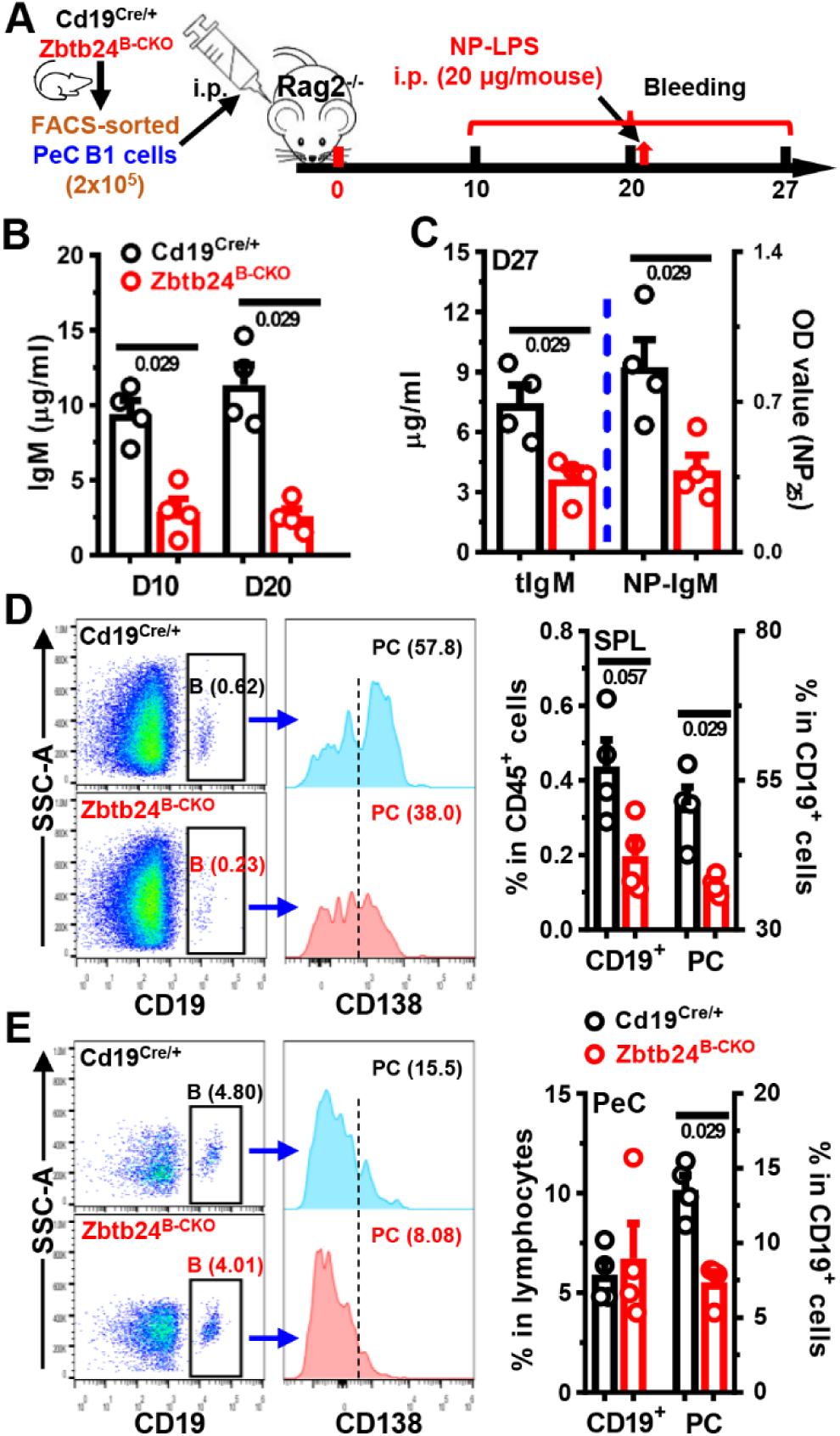
Impaired function of Zbtb24-deficient B1 cells *in vivo*. FACS-sorted peritoneal cavity CD19^+^B220^low^CD23^-^ B1 cells from control Cd19^Cre/+^ and Zbtb24^B-CKO^ mice (female, 12 weeks of age) were adoptively transferred into Rag2^-/-^ recipient mice (female, 8 weeks of age) intraperitoneally (2×10^5^ cells/200 μl PBS/mouse). On D20 post injection, Rag2^-/-^ recipient mice were immunized with NP-LPS (20 μg/200 μl PBS/mouse) intraperitoneally. Blood was taken at indicated times and cells in peritoneal cavities (PeC) & spleens (SPL) were analyzed by flow cytometry on D27. **A**, a schematic flow-chart showing the experiment setup. **B&C**, bar graphs showing levels of total (**B,** and tIgM in **C)** & NP-specific (NP-IgM in **C)** IgM in sera of recipient mice on D10 & D20 (**B**) or D27 (**C**). **D&E**, representative offset histograms/bar graphs showing the percentages of CD19+ B cells & CD19^+^CD138^+^ plasma cells (PC) within gated B cells in the spleens (SPL, **D**) and peritoneal cavities (PeC, **E**) of recipients implanted with Cd19^Cre/+^ or Zbtb24^B-CKO^ B1 cells. Each dot represents a single recipient mouse, and numbers below horizontal lines indicate *P* values.

Upon activation, some peritoneal B1 cells transit to the spleen to divide and differentiate into PCs, while others stay and rapidly differentiate into PCs even without division (Yang et al., 2007). We thus compared the number of B & PCs in the spleens and PeC of recipients. The percentages of Zbtb24-deficient CD19^+^ B cells and CD138^+^ PCs were significantly reduced in spleens, while only the latter was decreased in the PeC of recipients (**Figure 4D&E**).

In sum, our data (depicted in **Figures 3&4**, and **S6&7)** indicate that lack of Zbtb24 suppresses the differentiation of B1 cells toward PCs and thereby reduces the antibody levels against TID-Ags *in vivo*. Given that surface levels of CD11b, which couples with CD18 to form Mac-1/CR3 and governs the migration of B1 cells from the PeC into the spleens (Franke et al., 2020; Yang et al., 2007), and surface IgM were comparable on Zbtb24-deficient and -sufficient peritoneal B1 cells (**Figure S8A&B**), it is unlikely that reduced PC differentiation was attributable to impaired migration or BCR-triggering.

### Zbtb24-deficiency impedes the PC differentiation of B1 cells without compromising their survival, activation and proliferation *in vitro*

Upon activation by LPS *in vitro*, the differentiation of purified peritoneal B1, but not B2, cells towards CD138^+^ PCs was significantly impaired by Zbtb24 depletion (**Figure 5A-C**). Accordingly, levels of IgM & IgG3 in culture supernatants were reduced in LPS-cultivated Zbtb24-null B1 cells (**Figure 5D**). Given that B1 cells are more sensitive to TLR agonists-induced PC differentiation than FOB cells (Genestier et al., 2007), we further cultured splenic B cells with anti-IgM/CD40 or high amounts of LPS to mimic the T-cell dependent or independent stimulations, respectively. Again, deficiency of Zbtb24 did not suppress the PC differentiation of splenic B2 cells as well as their antibody producing abilities in these conditions (**Figure S8C-E**), corroborating our *in vivo* findings depicted in **Figures 2D&G** and **S4D**. Zbtb24-deficiency did not reduce the viabilities and cell numbers in these B1/B2 cultures (**Figures 5C**, and **S9A, D&E**). Moreover, the ability of B1/B2 cells to upregulate surface activation makers (CD69 & CD86) were not impaired by Zbtb24 depletion, excluding a generally disrupted signaling transduction downstream of TLR4 in B cells deprived of Zbtb24 (**Figure S9B, C&F**).

**Figure 5.**
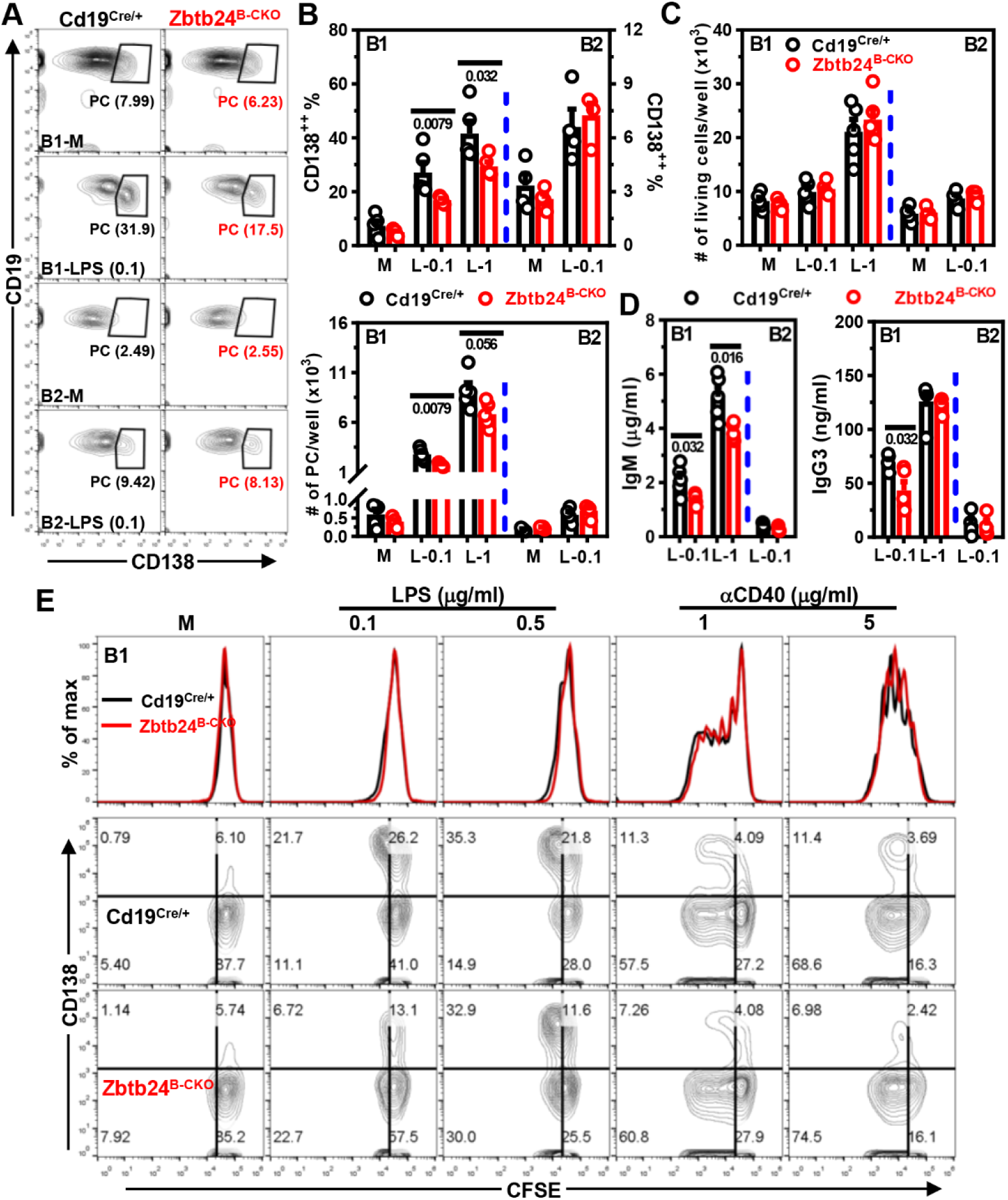
Zbtb24-deficiency specifically inhibits LPS-induced differentiation of peritoneal B1 cells toward PCs *in vitro*. CD19^+^B220^low^CD23^-^ B1 and CD19^+^B220^high^CD23^+^ B2 cells were FACS-sorted from peritoneal cavities of Cd19^Cre/+^ & Zbtb24^B-CKO^ mice, and subsequently cultured (2-5 x 10^4^ cells/well in 96-U bottom plate) in medium (M), 0.1/1 μg/ml LPS (L-0.1/L-1, respectively) for 3 days. **A**, representative contour-plots showing the percentages of CD19^low^CD138^+^ PCs in differentially cultured B1 or B2 cells on day 3. **B&C**, bar graphs showing the percentages/absolute numbers of PCs (**B**) or total living cells (**C**) 3 days post stimulation. **D**, bar graphs showing the IgM and IgG3 levels in culture supernatants of LPS-stimulated B1 or B2 cells on day 3. Antibody levels in supernatants of B cells cultured in medium were too low to be detected after 5x dilution. Each dot represents a single mouse of the indicated genotype (male, 10 weeks of age). **E**, representative overlayed histograms showing the comparable division profiles (*top panel*) or contour-plots showing the proliferation (CFSE) *vs.* differentiation (CD138, *middle and bottom panels*) of cultured peritoneal B1 cells isolated from Cd19^Cre/+^ *vs.* Zbtb24^B-CKO^ mice (male, 14 weeks old, n=3/4 for Cd19^Cre/+^ and Zbtb24^B-CKO^, respectively). B1 cells were labelled with the cell division tracker CFSE (10 μM) and subsequently cultured in medium (M), LPS (0.1/0.5 μg/ml) or anti-IgM (αCD40, 1/5 μg/ml) for 3 days before flow cytometry analysis. Numbers below horizontal lines in **B-D** indicate *P* values.

As reported previously (Sindhava and Bondada, 2012), B1 cells were anergic to BCR triggering and thus failed to undergo activation upon anti-IgM stimulation *in vitro* (**Figure S9B, C&F**). To further determine the role of Zbtb24 in cell proliferation, we labelled B1 cells with cell division tracker CFSE before culture. Interestingly, LPS stimulation induced massive PC differentiation accompanied by minimal cell divisions, whereas anti-CD40 elicited robust proliferation concurrent with little CD138 upregulation (**Figure 5E**), implying that B1 cells were able to uncouple PC differentiation from the cell division process. Of note, Zbtb24-depletion significantly repressed PC differentiation of B1 cells without impeding their mitoses induced by LPS or anti-CD40 *in vitro* (**Figure 5E**).

Based on these data, we conclude that Zbtb24 promotes the PC differentiation of B1 cells independent of their survival, activation and proliferation *in vitro*.

### Zbtb24-deficiency inhibits the PC differentiation of B1 cells *via* attenuating heme synthesis

We next performed RNA-Seq analysis to explore how Zbtb24 regulates the differentiation of B1 cells. Upon activation by LPS, Zbtb24-deficient peritoneal B1, but not splenic B2 cells, expressed significantly lower levels of hallmark genes involved in PC differentiation, UPR (unfolded protein response) and protein secretion, such as *Prdm1* (the driving force of PC differentiation), *Sd1* (encoding CD138), *Irf4* and *Xbp1* (**Figure 6A, B&D**, and **Table S1**). Of note, expressions of Bach2 & Bcl6, two major TFs repressing Prdm1, were not influenced by Zbtb24 depletion in B1 cells (**Figure 6B**). Therefore, it is unlikely that the B1-specific effect of Zbtb24 on PC differentiation is attributable to its direct binding and regulation of these key regulators.

**Figure 6.**
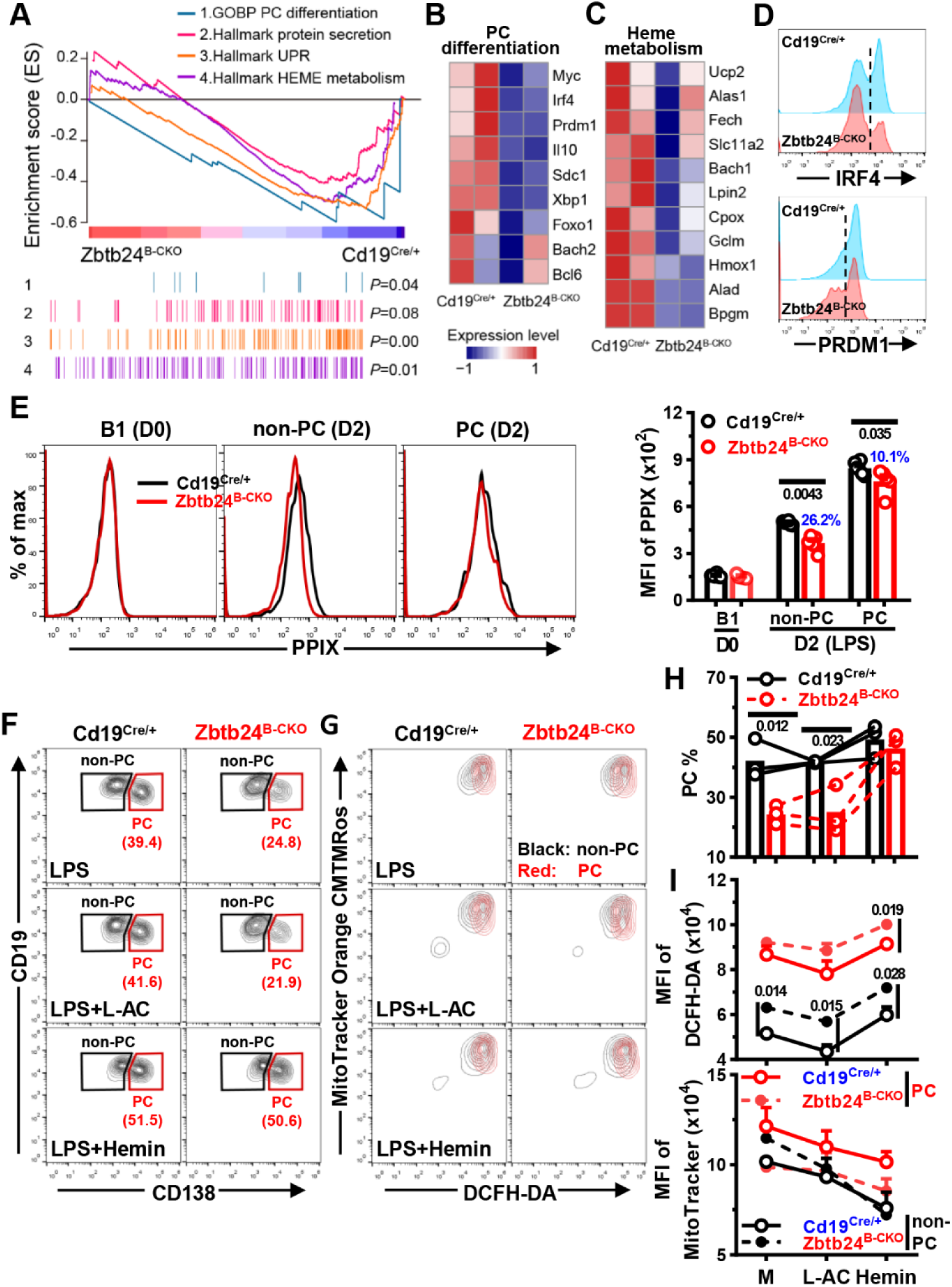
Zbtb24-deficiency blunts the differentiation of B1 cells into PCs through repressing the biosynthesis of heme. FACS-sorted peritoneal CD19^+^B220^low^CD23^-^ B1 cells were stimulated with 0.1 μg/ml LPS in 96-U bottom plate for 24 hrs (RNA-seq, **A-C)** or 2-3 days without/with additional L-AC (250 μM)/Hemin (25 μM) to induce PC differentiation (**D-I**). **A**, GSEA plots of genes involved in PC differentiation, protein secretion, unfolded protein response (UPR), and heme metabolism in LPS-stimulated Zbtb24^B-CKO^ *vs.* Cd19^Cre/+^ B1 cells. **B&C**, heatmaps showing the dysregulated genes regulating the differentiation of PC cells (**B**) or heme metabolism in cells (**C**) identified by RNA-seq analysis. The complete lists of enriched genes regulating UPR and heme metabolism were provided in Figure S10B. **D**, representative half-offset histograms showing the levels of intracellular IRF4 & PRDM1 on in LPS-stimulated B1 cells on D2. **E**, representative overlayed histograms (*left*) and cumulative data (*right*) showing the levels of intracellular PPIX in resting control *vs.* Zbtb24^B-CKO^ B1 cells (D0) or gated CD138^-^ non-PC & CD138^+^ PC cells in LPS-stimulated cultures on D2. **F&H**, representative contour-plots (**F**) and bar graphs (**H**) showing the percentages of CD19^low^CD138^+^ PCs in differentially cultured Cd19^Cre/+^ *vs.* Zbtb24^B-CKO^ B1 cells on day 3. **G**, representative overlayed contour-plots showing the intracellular ROS levels (visualized by DCFH-DA) and mitochondrial mass/membrane potentials (detected by MitoTracker Orange CMTMRos) in gated CD138^-^ non-PC (black) *vs.* CD138^+^ PC (red) cells in cultured B1 cells on day 3. **I**, bar graphs showing the ROS levels (MFI of DCFH-DA, top) or mitochondrial mass/membrane potential (MFI of MitoTracker, bottom) in gated non-PC/PC cells in Cd19^Cre/+^ *vs.* Zbtb24^B-CKO^ B1 cultures. Gates for CD138^-^ non-PC and CD138^+^ PC were illustrated in **F**. Each dot in **E&H** represents a single mouse of the indicated genotype (male, 12-14 weeks of age), while results in **I** are expressed as mean ± SEM (n=3). Numbers in **E**, **H &I** indicate *P* values determined by student *t*-test. Blue numbers in **E** denote percents of reduction.

Enormous studies recently showed that the activation and function of lymphocytes are intertwined with metabolic reprogramming & adaption (Boothby and Rickert, 2017; Foh et al., 2022; Jellusova, 2018). In association with the altered function, multiple genes relating to nutrients uptake and metabolism, such as Slc2a1/Glut1, Slc16a3/Mct3, Ak4, Pdk1, Pfkl, and Tpi, were significantly upregulated in Zbtb24^B-CKO^ B1 cells (**Figure S10A**). Accordingly, GSEA identified a positive enrichment of gene sets regulating the glycolysis and pyruvate metabolism in Zbtb24-null B1 cells (**Figure S10C**). By contrast, a panel of genes involved in heme metabolism were downregulated in Zbtb24^B-CKO^ cells (**Figures 6A&C**, and **S10B**), albeit that the differences did not reach the cutoff value. Expressions of Alad (5-aminolevulinic acid dehydratase) and Cpox (Coproporphyrinogen oxidase), two enzymes that directly catalyze reactions along the heme synthesis pathway in cytosol and mitochondrial intermembrane space, respectively (Ajioka et al., 2006), were consistently reduced in Zbtb24-null B1 but not B2 cells (**Figure 6C**, and **Table S1**). Moreover, the level of Hmox1, a heme-induced oxygenase that mediates its degradation and serves as a readout for intracellular heme content (Watanabe-Matsui et al., 2011), was mildly decreased in Zbtb24^B-CKO^ B1 cells as well (**Figure 6C**). Given that heme promotes Prdm1 expression and PC differentiation of B cells by inactivating Bach2 (Watanabe-Matsui et al., 2011), we reasoned that heme biosynthesis may be attenuated in Zbtb24-depleted B1 cells, thereby leading to impaired PC differentiation.

Indeed, Zbtb24-depletion significantly mitigated LPS-induced upregulation of protoporphyrin IX (PPIX), the final substrate of heme biosynthesis (Ajioka et al., 2006), in B1 cells, albeit no differences were observed in resting cells (**Figure 6E**). Notably, ablation of Zbtb24 resulted in a more pronounced PPIX reduction in CD138^-^ non-PC cells than that in CD138^+^ PC cells (26.2% *vs.* 10.1%, **Figure 6E**). Moreover, addition of exogenous hemin in cultures almost completely abrogated the differentiation defects of Zbtb24-deficient B1 cells (**Figure 7F&H**), while its supplementation similarly promoted/suppressed PC differentiation/CSR in control *vs.* Zbtb24^B-CKO^ splenic B cells (**Figure S11A&B)**. Together, these data indicate that Zbtb24 specifically promotes the PC differentiation of B1 cells *via* augmenting heme synthesis. Exogenous hemin only slightly potentiated the PC differentiation of control B1 cells (**Figure 7F&H**), implying that activated peritoneal B1 cells may contain abundant intracellular heme to facilitate their accelerated and heightened PC differentiation.

**Figure 7.**
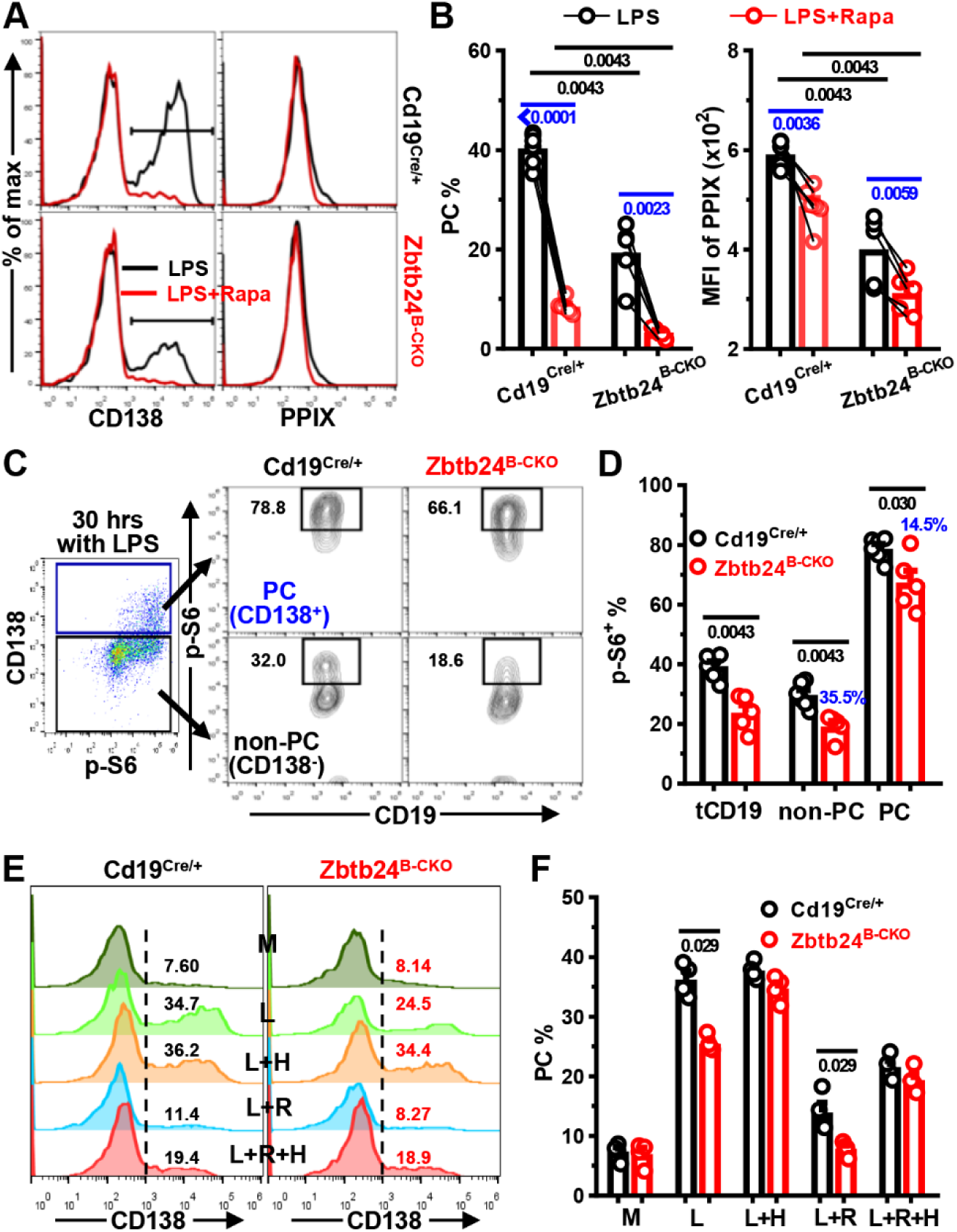
Zbtb24-ablation represses the mTORC1 activity, which promotes heme synthesis and PC differentiation of B1 cells. Peritoneal CD19^+^B220^low^CD23^-^ B1 cells were stimulated with 0.1 μg/ml LPS in the absence/presence of Rapamycin (Rapa, 10 nM) up for 2 days (**A-D**). **A&B**, representative overlayed histograms (**A**) and bar graphs (**B**) showing the PC differentiation (*left panel*) and PPIX expression (*right panel*) in B1 cells cultured with LPS ± Rapa on day 2. **C&D**, representative FACS-plots (**C**) and bar graphs (**D**) showing the expression of phosphorylated ribosomal protein S6 (p-S6) in gated CD138^+^ PC or CD138^-^ non-PC cells derived from Cd19^Cre/+^/Zbtb24^B-CKO^ B1 cells after stimulation with LPS for 30 hrs. tCD19 denotes total CD19^+^ B cells and blue numbers indicate percent of reduction in gated Zbtb24^B-CKO^ populations compared to the corresponding control counterparts (**D)**. **E&F**, representative histograms (**E**) and bar graphs (**F**) showing the PC differentiation of B1 cells cultured in medium (M), LPS (L), and LPS plus Hemin (25 μM, L+H), Rapamycin (10 nM, L+R) or both (L+R+H) for 2 days. Each dot represents a single mouse of the indicated genotype (12-week males in **A-D**, and 9-week females in **E&F**). Black/blue numbers below horizonal lines indicate *P* values determined by Mann-Whitney test or paired *t*-test, respectively.

### Zbtb24-depletion represses the heme synthesis partially through mTORC1 in B1 cells

It has been recently shown that endogenous ROS inhibits heme synthesis, and activated splenic B cells with intermediate mitochondrial mass/membrane potential are predisposed to become PCs (Jang et al., 2015). We thus wondered whether Zbtb24 promoted heme synthesis in B1 cells *via* regulating intracellular ROS and mitochondrial function. Compared to activated splenic B cells, B1 cells had significantly higher levels of ROS, which were further increased by Zbtb24-depletion (**Figures 7F-I**, and **S11C&D**). However, addition of the antioxidant L-ascorbic acid (L-AC) failed to promote PC differentiation in peritoneal B1 cells, albeit that it did reduce ROS levels in B1 cell and augment the PC differentiation of splenic B2 cells (**Figures 7F-I**, and **S11A&B**). The mitochondrial mass/membrane potential was markedly lower in splenic B2-derived PCs as previously reported (Jang et al., 2015), but no such differences were observed in B1 cultures, and no significant impact of Zbtb24-deficiency on mitochondrial mass/membrane potential of B cells was observed (**Figures 7G&I**, and **S11C&D**). Thus, the metabolic reprogramming and mitochondrial function differ significantly between B1 *vs.* B2 cells along their differentiation toward PCs. Because ROS represses the final step of heme synthesis by inhibiting the addition of ferrous ions to PPIX (Jang et al., 2015), it is unlikely that disturbed ROS levels were unlikely responsible for the attenuated PPIX accumulation in Zbtb24-deficient B1 cells.

Expressions of Slc3a2 & Slc7a5, encoding the respective heavy & light chain of CD98, positively correlate with mTORC1 activity in B cells (Tsui et al., 2018). RNA-Seq data showed that levels of Slc3a2 & Slc7a5 were decreased, coinciding with the disturbed mTORC1 signaling in activated Zbtb24-null B1 cells as revealed by GSEA (**Figure S10A&C**). mTORC1 activity regulates the protein synthesis & metabolism of mammalian cells, and promotes PC differentiation of murine splenic B cells partially *via* augmenting heme synthesis (Tsui et al., 2018). In keeping with these findings obtained in B2 cells, inclusion of an mTORC1 specific antagonist, Rapamycin, significantly suppressed the PC differentiation and PPIX levels in peritoneal B1 cells (**Figure 7A&B**), demonstrating that mTORC1 promotes heme synthesis and PC differentiation of B1 cells as well. In line with this notion, Zbtb24-depletion significantly repressed the expression of phosphorylated ribosomal protein S6 (p-S6), an event controlled by mTORC1 activity (Tsui et al., 2018), in LPS-stimulated B1 cells (**Figure 7C&D**). Notably, the inhibition was more pronounced, akin to PPIX contents, in CD138^-^ cells as compared to that in CD138^+^ PCs (**Figures 6E**, and **7C&D**), indicating that the impaired mTORC1 activity underlie, at least partially, the attenuated heme synthesis and PC differentiation of Zbtb24-null B1 cells.

We noticed that PC differentiation and intracellular PPIX levels were still considerably reduced in mTORC1 repressed (i.e., cultured with LPS plus Rapamycin) Zbtb24-null B1 cells (**Figure 7A&B**). To determine the extent to which the two factors (mTORC1 activity *vs.* heme accumulation) contribute to the defected PC differentiation of Zbtb24-deficient B1 cells, we differentiated B1 cells in the presence of Rapamycin/hemin alone or both (**Figure 7E&F**). Supplementation of hemin not only completely reverted the defects caused by Zbtb24-depletion as previously observed in **Figure 6F&H**, but also partially relieved the inhibitory effect of Rapamycin on PC differentiation of B1 cells. Of note, control and Zbtb24-null B1 cells exhibited comparable PC differentiation in the presence of both Rapamycin and hemin (**Figure 7E&F**), implying that attenuated intracellular heme synthesis is mainly responsible for the differentiation defects of Zbtb24-deficient B1 cells.

Collectively, our data show that Zbtb24 promotes the PC differentiation of B1 cells mainly through augmenting heme synthesis, and the attenuated heme biosynthesis in Zbtb24-depleted B1 cells is partially attributed to their impaired mTORC1 activity.

### No effect of Zbtb24-deficiency on PC differentiation of MZB cells

Akin to B1 cells, MZB cells are specialized in responses to TID-Ags and more sensitive, compared to conventional FOB cells, to TLR agonists-induced proliferation (**Figure 8A**) and PC differentiation (Genestier et al., 2007). We thus questioned whether Zbtb24 exerts a similar function in this innate-like B cell subset as well. LPS stimulation induced robust cell division and PC differentiation of MZB cells, while BCR-crosslinking elicited massive proliferation accompanied by mild PC generation (**Figure 8A-C**). Notably, neither the proliferation nor the PC differentiation of LPS-cultivated MZB cells were affected by Zbtb24 depletion, and the addition of exogenous hemin enhanced PC differentiation of Zbtb24 deficient or sufficient MZB cells to the same extent (**Figure 8A-D**). Hence, depletion of Zbtb24 does not impact the PC differentiation of MZB cells.

**Figure 8.**
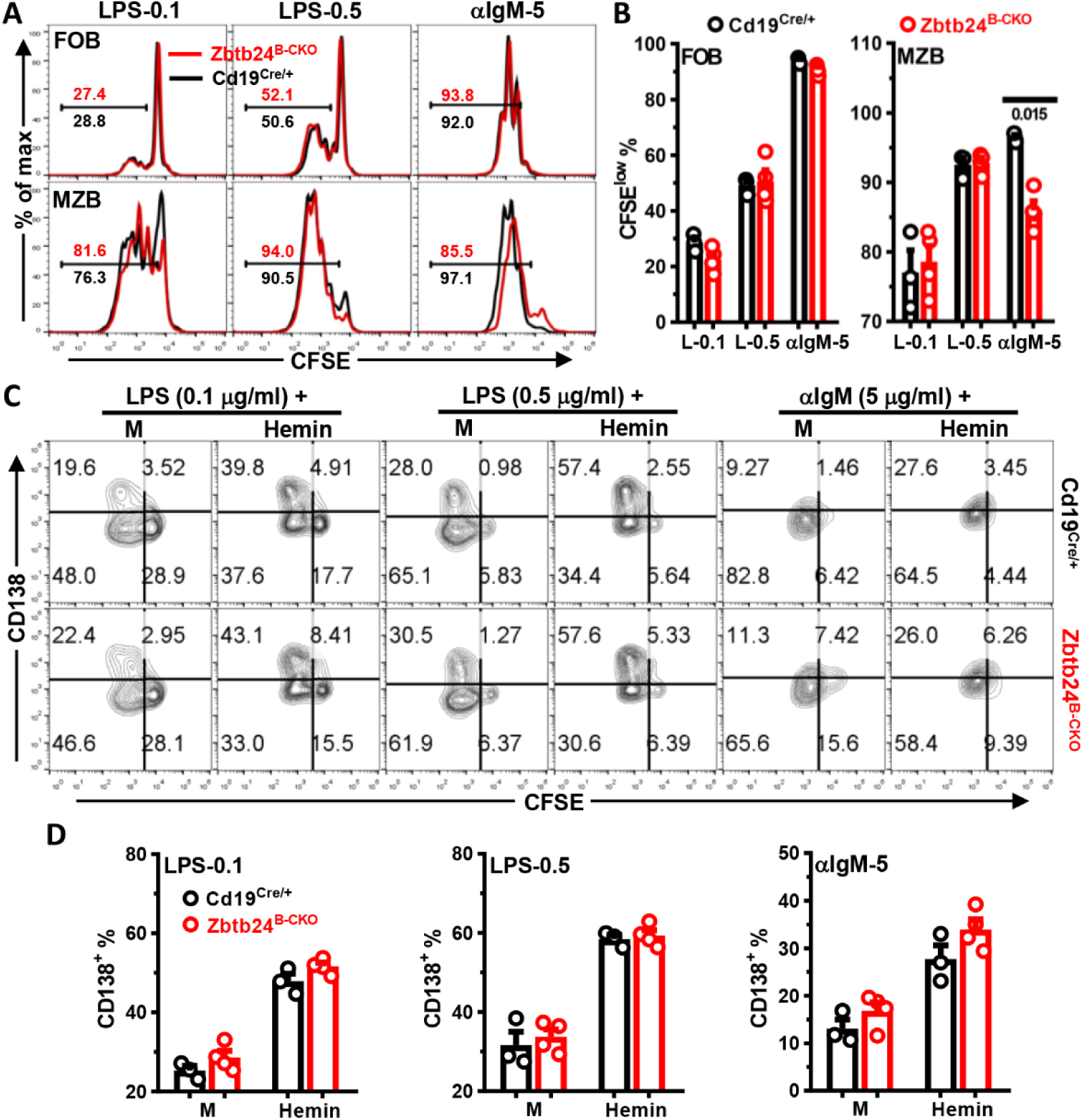
Zbtb24-deficiency has no effect on the differentiation, but slightly reduces BCR-triggered proliferation of MZB cells *in vitro*. CD19^+^CD23^high^CD21^low^ FOB and CD19^+^CD23^low^CD21^high^ MZB cells were FACS-sorted from spleens of Cd19^Cre/+^ & Zbtb24^B-CKO^ mice (male, 14 weeks old), labeled with CFSE (10 μM), and then cultured (∼1 x 10^5^ cells/well in 96-U bottom plate) with LPS (0.1/0.5 μg/ml), αIgM (5 μg/ml) in the absence/presence of exogenous Hemin (25 μM). On day 3, cells were collected, stained with anti-CD138 before flow cytometry analysis. **A**, representative overlayed histograms showing the cell division profiles (CFSE) of cultured control *vs.* zbtb24^B-CKO^ FOB (top) and MZB (bottom) cells. Numbers indicate the percentages of divided CFSE^low^ cells. **B**, bar graphs showing the percentages of divided CFSE^low^ cells in cultured FOB and MZB cells. **C**, representative contour-plots showing the proliferation (CFSE) *vs.* differentiation (CD138) of splenic control (top) *vs.* zbtb24^B-CKO^ (bottom) MZB cells in different cultures. **D**, bar graphs showing the percentages CD138^+^ PC cells in differently stimulated MZB cells purified from control *vs.* zbtb24^B-CKO^ mice. Each symbol represents a single mouse of the indicated genotype, and numbers below horizonal lines in **B** indicate *P* values determined by student *t*-test.

Notably, in stark contrast to the massive PC differentiation but little proliferation of peritoneal B1 cells (**Figure 5E**), the vigorous LPS-elicited differentiation of MZB cells was companied by extensive cell division (**Figure 8A-C**). Moreover, depletion of Zbtb24 resulted in a subtle but nevertheless consistent decrease of divided CFSE^low^ MZB cells in anti-IgM-stimulated cultures (**Figure 8A&B**). This slightly reduced BCR-elicited proliferations of MZB cells may contribute to the decreased antibody levels in Zbtb24^B-CKO^ mice after TID-Ag immunization as well. We speculated that the distinct metabolic adaptation and energy/substrate allocation between proliferation and differentiation in B1 *vs.* MZB cells, triggered by different stimuli, might underlie the distinct effects of Zbtb24 in these cells.

## DISCUSSION

The clinical and immunological presentations of *ZBTB24*-disrupted ICF2 patients imply a prominent role of ZBTB24 in the terminal differentiation and antibody secretion of human B lymphocytes *in vivo* (Banday et al., 2020; Weemaes et al., 2013). Thus, we here systemically characterized the cell autonomous role of Zbtb24 in B cell development, differentiation and function by using the conditional knockout mice. Unexpectedly, our data demonstrate that B-cell Zbtb24 is virtually dispensable for TD-Ag-induced GC reactions & antibody responses *in vivo*, but its deficiency specifically impeding the PC differentiation B1 cells and thereby reduces TID-Ags-induced antibody productions. Our study thus not only provides a plausible explanation for the recurrent respiratory and gastrointestinal infections in ICF2 patients, but also is relevant for B1 cell-mediated barrier defenses and immune regulations.

Despite the reduced levels of serum antibodies and peripheral CD19^+^CD27^+^ Bm cells, total amounts of circulating T and B lymphocytes are normal in most ICF cases (Banday et al., 2020; Weemaes et al., 2013). Moreover, detailed anatomical and immunological examinations revealed the absence of GC structure in one ICF2 patient devoid of Zbtb24, and downregulating its expression inhibits the proliferation of a human B cell line by indirectly repressing Prdm1 & Irf4 *in vitro* (Kloeckener-Gruissem et al., 2005; Liang et al., 2016; Weemaes et al., 2013). Given the essential role of GC in the propagation of Bm and antibody-secreting PCs (Victora and Nussenzweig, 2022), these findings strongly indicate that ZBTB24 controls *in-vivo* antibody responses through regulating the GC structure/reaction rather than the general development/maintenance of lymphocytes in humans. In partial agreement of this notion, specific deletion of Zbtb24 in B cell compartment from the CD19^+^ pro-B stage onwards (*Cd19*-Cre) has no effect on the later development, maintenance and phenotype of B cells in the BM & periphery of mice. However, contrary to the Bm and antibody defects in *ZBTB24*-disrupted ICF2 patients, neither the GCB numbers, nor the primary/secondary antibody responses elicited by TD-Ags are significantly decreased in Zbtb24^B-CKO^ mice, regardless of antigen doses and adjuvant types used for immunization. In keeping with these *in-vivo* findings, Zbtb24-depletion does not impair the *in-vitro* survival, activation, proliferation and differentiation of FOB cells, the main responders to TD-Ags. Furthermore, mice with a specific depletion of Zbtb24 in T cells (*Cd4*-Cre) or the whole hematopoietic lineages (*Vav*-Cre) exhibit normal B cell development and largely intact TD-Ag-induced antibody responses *in vivo* as well (unpublished data).

Hence, hematopoietic Zbtb24 is dispensable for B cell development/maintenance and the canonical TD-Ag-induced antibody responses *via* GC reactions in mice. The latter is clearly distinct from *ZBTB24*-disrupted ICF2 patients with hypo- or a-gammaglobulinemia and lack of circulating Bm cells (Banday et al., 2020; Weemaes et al., 2013). The discrepancies in (memory) humoral responses between ICF2 patients and aforementioned Zbtb24-deficient mice may stem from the distinct functions of Zbtb24 across species as evidenced by the fact that Zbtb24-null embryo is only viable in humans but not in mice (Wu et al., 2016). It is also possible that Zbtb24 may regulate TD-Ag-induced GC reaction and antibody responses beyond the hematopoietic system, for instance, through supporting the formation of proper B cell follicular cell networks that steers the GC output (Lutge et al., 2023). Nonetheless, a functional substitution of Zbtb24 by other Zbtb proteins, albeit unlikely based on our RNA-Seq data in splenic B cells, could not be completely excluded.

By contrast, our data highlight an important role of B-cell Zbtb24 in humoral responses against TID-Ags *in vivo.* Upon immunization with a type I or type II TID-Ag, Zbtb24^B-CKO^ mice generate significantly less antibody-secreting PCs in the presence of normal amounts of B1, FOB and MZB cells. At the cellular level, depletion of Zbtb24 markedly restrains the ability of B1 cells to differentiate into PCs and produce antibodies both *in vitro* upon LPS stimulation and *in vivo* after transfer into Rag2^-/-^ recipients without impairing their *in-vitro* survival, activation and proliferation. MZB cells are the other main responders, beyond B1 cells, to TID-Ags, and these two innate-like B cell subsets behave similarly in many aspects. For example, both B1 and MZB are hypersensitive to TLR agonists-induced PC differentiation (Genestier et al., 2007). Strikingly, Zbtb24-ablation does not markedly impact the proliferation & PC differentiation of MZB cells. Therefore, our study discloses that B1 cells are the main target B-cell subset through which Zbtb24 augments TID-Ag-elicited antibody productions in *vivo*.

Unlike conventional FOB cells that need to undergo extensive proliferation before terminal differentiation, B1 cells do no respond readily to BCR-induced clonal expansion, possibly owing to the highly expressed inhibitory co-receptors like CD5 & Lyn. Instead, B1 cells secrete natural IgM at steady state in the absence of antigenic stimulation, and upon encountering with non-specific inflammatory/pathogen-associated stimuli, they promptly and vigorously differentiate into antibody (mainly IgM)-secreting cells (Genestier et al., 2007; Holodick and Rothstein, 2013; Sindhava and Bondada, 2012). These polyreactive IgM not only participates in the maintenance of tissue homeostasis and prevention of autoimmune reactions, but also constitutes an important first-line defense against infections before the activation of adaptive immune system by facilitating the clearance of autologous dead/dying cells or invading pathogens, respectively (Morris et al., 2019). Moreover, B1 cells may regulate the immune response independent of antibody secretion. Both at steady state and after activation, B1 cells may function as a sort of regulatory B (Breg) cells by producing copious amounts of anti-inflammatory cytokine IL-10 (Catalan et al., 2021). Zbtb24-deficeint B1 cells produce less IgM and IL-10, it is thus tempting to speculate that the dysfunctional B1 cells contribute to the commonly-occurred infections as well as the uncommonly-observed autoimmune manifestations in ICF2 patients (Banday et al., 2020), albeit that the putative surface markers defining human B1 cells remain controversial so far (Sanz et al., 2019).

The promptly upregulated PC differentiation and antibody-producing ability of B1 cells upon activation imply that they are transcriptionally and metabolically well prepared for the high secretory capacity. Indeed, innate-like B1 and MZB cells constitutively express more Prdm1 while less Bcl6, and are bioenergetically more active than conventional FOB cells (Genestier et al., 2007; Jellusova, 2018). Moreover, sustained mTORC1 signaling coordinates an immediate UPR-affiliated transcriptome profile preceding the expression of Xbp1, and confers MZB cells the ability to undergo rapid and robust PC differentiation even in the presence of cell cycle inhibitors (Gaudette et al., 2020; Gaudette et al., 2021), which mirrors the unique mitosis-independent PC differentiation of B1 cells as we showed here. In activated conventional splenic B cells, mTORC1 promotes heme synthesis partially by reducing intracellular ROS, and thereby augments their PC differentiation because heme induces Prdm1 by inactivating its repressor molecule Bach2 (Jang et al., 2015; Tsui et al., 2018; Watanabe-Matsui et al., 2011). Once synthesized, heme enhances mTORC1 activity, possibly through Prdm1 induction or modulating the mitochondrial function (Price et al., 2021; Tsui et al., 2018). We here showed that mTORC1 exerts a similar function in B1 cells as Rapamycin-mediated mTORC1 inhibition greatly inhibits the PC differentiation and heme accumulation in B1 cells as well. Given that Zbtb24-depletion reduces mTORC1 activity and PPIX content, and that hemin supplementation almost completely reverts the differentiation defects of Zbtb24-null B1 cells even in the presence of Rapamycin, our data reveal that Zbtb24 promotes the PC differentiation of B1 cells mainly *via* heme synthesis, which is partially mediated through mTORC1. Detailed mechanisms underlying the regulations of mTORC1 activity and heme synthesis in B1 cells by Zbtb24 merit further investigation.

Both human and murine Bm cells possess an enhanced heme signature than naive B cells, and supplementation of hemin promotes their differentiation towards PCs (Price et al., 2021). Thus, increased heme biosynthesis seems to be a conserved feature of B cells with accelerated & augmented PC differentiation potential, including B1, MZB & Bm cells. The comparable antibody levels between control and Zbtb24^B-CKO^ mice after secondary TD-Ag challenge indicate that Zbtb24 has little effect on heme metabolism and PC differentiation of murine Bm cells. Why Zbtb24 only influences the heme biosynthesis and PC differentiation of B1, but not other B cells, at least in mice, is intriguing. Although MZB and B1 cells are considered innate-like B cells and able to mount rapid antibody responses against invading pathogens before the adaptive immune responses are properly propagated and matured, these two cell types are functionally distinct at many aspects. For instance, our data demonstrate that only B1 cells are able to largely uncouple PC differentiation from cell division upon activation, albeit that MZB cells may differentiate into PCs in the presence of cell cycle blockers (Gaudette et al., 2021). It is thus conceivable that B1 cells, depending on the types of stimulation (i.e., LPS *vs.* anti-CD40 triggering), adopt distinct metabolic pathways to devote most of the anergy & biomolecule building blocks either to the PC differentiation/antibody secretion pathway or to the mitosis process. We hypothesize that in case of the PC differentiation pathway, B1 cells immensely and sharply elevate heme biosynthesis, and Zbtb24 somehow participates in this metabolic adaptation process partially by regulating mTORC1 activity. In MZB cells, this metabolic reprogramming is much milder and thus less affected by Zbtb24 deficiency. Despite that mice deficient in *Hmox1*, the heme degrading enzyme, have increased serum antibody levels, administration of hemin to mice does not augment antibody responses against TD- or TID-Ags (Kapturczak et al., 2004; Price et al., 2021; Watanabe-Matsui et al., 2011), indicating a complex effect of heme *in vivo*.

During the preparation of this manuscript, Chen’s group reported that ablation of Zbtb24 in B cell compartment (*Cd19*-Cre) had no effect on basal serum antibody levels and TD-Ag-induced GC reactions in mice as well (Ying et al., 2023), corroborating our findings in this aspect. They found that Zbtb24-depletion upregulated Il5ra expression, which results in augmented CD19 activity and impaired MZB proliferation induced by BCR (Ying et al., 2023). In partial agreement with their findings, BCR-triggered proliferation of MZB cells were impaired upon Zbtb24-depletion, albeit that the defect was far milder in our hands. Moreover, our RNA-Seq analysis in splenic B cells did not reveal any differences in Il5ra expression. These discrepancies may relate to different strategies used to generate the conditional knockout mice. Nonetheless, we here extend the role of Zbtb24 in humoral immune response by showing that lack of Zbtb24 significantly impedes the mTORC1 activity, heme synthesis and PC differentiation of B1 cells, leading to diminished TID-Ag-induced antibody productions *in vivo*. Thus, Zbtb24 may represent an important factor involved in regulating barrier defenses against pathogens through B1 cells.

## MATERIALS AND METHODS

### Mice

The *Zbtb24* conditional allele (*Zbtb24^loxp/+^*) was generated using the CRISPER/Cas9 strategy (Beijing CasGene Biotech. Co., Ltd). Briefly, C57BL/6 embryonic stem cells (ES) were transfected with the targeting vectors *via* electroporation. Correctly targeted clones, containing two loxp sites inserted into the flanking regions of *Zbtb24* exon 2 with the translation starting site (**Figure 1A**), were identified by PCR and injected into blastocytes to generate chimeric mice. Chimeric mice were then crossed with C57BL/6 wild type (WT) mice to obtain F1 mice with the germline-transmitted *Zbtb24^loxp/+^*allele. After multiple-step crossing the heterozygous *Zbtb24^loxp/+^*mouse with Cd19^Cre/+^ mouse on a C57BL/6 background (kindly provided by Prof. Biao Zheng, East China Normal University), mice with the specific deletion of *Zbtb24* in CD19^+^ B cells (Zbtb24^B-CKO^, *Cd19^Cre/+^Zbtb24^loxp/loxp^*) were generated. Primers used for ES clone selection and genotyping were listed in **Table S2**. To exclude the impact of *Cd19*-driven Cre expression on B cells (Zhao et al., 2022), littermate *Cd19^Cre/+^Zbtb24^+/+^* (Cd19^Cre/+^) mice were used as controls in all the analysis unless otherwise indicated. Immunodeficient *Rag2^-/-^* C57BL/6 mice with no mature T & B cells (Changzhou Cavens Laboratory Animal Co., Ltd) were used as recipients in adoptive transfer experiments. Mice were housed under specific pathogen-free (SPF) conditions and all experiments were performed in accordance with procedures approved by the Ethics Committee of Soochow University.

### Immunizations

Mice were immunized with the TD-Ag NP_19_-OVA emulsified in IFA (Sigma, 1:1) or adsorbed onto Imject Alum (Thermo Scientific, 1:1), or with the TID-Ag NP-AECM-Ficoll (NP-Ficoll)/NP-LPS intraperitoneally. NP_19_-OVA, NP-Ficoll and NP-LPS were all obtained from Biosearch Technologies.

### B-cell Isolation and Culture

BM cells were harvested by flushing the femurs of mouse, and peritoneal cells were isolated by lavaging the PeC with 10 ml PBS. Splenocytes were obtained by mechanically dissociating spleens in PBS, followed by passing through a 70 μM nylon mesh (BD Biosciences). Erythrocytes were removed by addition of ammonium chloride lysis buffer. Total CD19^+^ B, CD19^+^CD23^high^CD21^low^ FOB and CD19^+^CD23^low^CD21^high^ MZB cells in spleens, or peritoneal CD19^+^B220^low^CD23^-^ B1 and CD19^+^B220^high^CD23^+^ B2 cells were sorted out *via* a FACS Aria cell sorter III (BD Biosciences) with > 95% purity. MojoSort^TM^ Mouse Pan B Cell Isolation Kit II (Biolegend) were used to enrich CD19^+^ B cells in splenocytes before sorting.

Purified B cells were cultured in medium, LPS (L2630, Sigma), goat F(ab’)_2_ anti-mouse IgM (αIgM, SouthernBiotech) or rat anti-mouse CD40 (αCD40, 1C10, Biolegend) in the absence/presence of recombinant mouse IL-4, human TGF-β1 and B-cell activating factor (BAFF) (all from Peprotech) in 96 U-bottom plates at 37°C for indicated times. The complete culture medium was RPMI-1640 (Hyclone) supplemented with non-essential amino acids (Sigma), 10% heat-inactivated fetal bovine serum (FBS, Gibco), 100 U/ml penicillin (Beyotime), 100 μg/ml streptomycin (Beyotime), 50 μM β-mercaptoethanol (Sigma), 1 mM sodium pyruvate and 10 mM HEPES (Hyclone). In some cases, L-ascorbic acid (L-AC, #A92902, Sigma), hemin (#9039, Sigma), Rapamycin (Absin), and corresponding vehicle controls were added at indicated concentrations from the beginning of culture.

### Adoptive Transfer Experiments

CD4^+^ T cells were isolated with MojoSort^TM^ Mouse CD4 T cell isolation kit (Biolegend) from spleens of WT mice with > 90% purity. Purified splenic CD19^+^ B cells, from Cd19^Cre/+^ or Zbtb24^B-CKO^ mice, were mixed with CD4^+^ T cells at the ratio of 2:1 before being injected into Rag2^-/-^ recipients *via* the tail vein (5 x 10^6^ B cells plus 2.5 x 10^6^ CD4^+^ T cells/200 μl PBS/mouse). One day later, recipient mice were immunized with NP_19_-OVA emulsified in IFA. To analyze the function of B1 cells *in vivo*, FACS-sorted peritoneal B1 cells were injected into Rag2^-/-^ recipients intraperitoneally (2 x 10^5^ cells/200 μl PBS/mouse) followed by NP-LPS immunization.

### ELISA

Mouse sera or B-cell culture supernatants were collected at indicated time points and immediately stored at -20℃ until the analysis for antibody levels by ELISA. In brief, ELISA plates (Nunc) were coated with goat anti-mouse Ig (SouthernBiotech, 1010-01, 1:1000) or NP_25_-BSA (Biosearch Technologies, 5 μg/ml) to capture all murine Igs or NP-specific antibodies, respectively. After washing with PBS containing 0.05% Tween-20, wells were blocked with PBS containing 3% BSA before incubation with diluted sera or culture supernatants. The optimal dilutions for each Ig subtype were determined by preliminary experiments. Total or NP-specific IgM, IgG1, IgG2b, IgG2c, IgG3 and IgA levels were detected by using HRP-coupled goat anti-mouse subtype-specific secondary antibodies (SouthernBiotech).

### Western Blot and Q-PCR Analysis

Cells were lysed for 1 hr on ice in RIPA lysis buffer (Beyotime) supplemented with protease and phosphatase inhibitors (Selleck). Proteins were separated in an SDS-PAGE 10% gel and subsequently blotted onto a PVDF membrane (Millipore). After incubation with blocking buffer (Tris-buffered saline with 5% nonfat milk), the membrane was incubated with primary antibodies overnight at 4°C, followed by a further incubation with secondary antibodies at room temperature (RT) for 2 hrs. Signals were visualized with the Tanon 3500 Gel Imaging System (Tanon). The following antibodies were used: anti-Zbtb24 (PM085, MBL Life Science, 1:1000), anti-Gapdh (AF0006, Beyotime, 1:1000), HRP-coupled donkey anti-Guinea pig (H+L, 706-035-148, Jackson ImmunoResearch Lab) and HRP-conjugated horse anti-mouse IgG (7076S, CST).

Total RNA was extracted from indicated cells by the Total RNA Kit II (OMEGA). cDNA was subsequently synthesized using Reverse Transcriptase M-MLV (TakaRa) according to manufacturer’s instructions. mRNA levels of indicated genes were quantitatively determined by SYBR-green technology on an ABI-StepOnePlus Sequence Detection System (Applied Biosystems). Sequence of primers used for RT-qPCR analysis were listed in **Table S2**.

### Flow Cytometric Analysis

Single cell suspensions were prepared and surface molecules were stained at 4°C for 30 mins with optimal dilutions of each antibody. The following antibodies were used: anti-mouse B220 (RA3-6B2), CD45 (30-F11), CD19 (6D5), CD23 (B3B4), CD5 (53-7.3), CD38 (90), CXCR4 (L276F12), CD69 (H1.2F3), CD86 (PO3), IgG1 (RMG1-1) and IgG2b (RMG2b-1) (all from Biolegend); anti-mouse CD21/35 (eBio4E3 or eBio8D9), CD43 (eBioR2/60), CD93 (AA4.1) and IgM (II/41) (all from eBioscience); and anti-mouse CD138 (281-2), CD95 (Jo2) and IgG3 (R40-82) (all from BD Biosciences). Sometimes 7-AAD (Biolegend) and NP-Ficoll-FITC (NP-FITC, Biosearch Technologies) were additionally used to visualize NP-specific B cells as previously reported (Zhao et al., 2022). After staining, cells were washed twice with PBS, suspended in 300 μl PBS, and fixed volumes of cells were processed with the Attune® NxT Acoustic Focusing Cytometer (Thermo Scientific). Data were analyzed by FlowJo software (BD Biosciences).

To measure the mitochondrial membrane potential and intracellular ROS, cells were incubated in pre-warmed RPMI-1640 containing MitoTracker^TM^ Orange CMTMRos (Thermo Scientific, 50 nM) and DCFH-DA (Beyotime, 5 μM) at 37°C for 30 mins. After extensive washing, cells were further stained with antibodies against surface molecules as indicated above.

For intracellular detection of TFs, cells were fixed/permeabilized with Foxp3 staining buffer set (eBioscience) before incubation with antibodies against BCL6 (7D1), PRDM1 (5E7) and IRF4 (IRF4.3E4) (all from Biolegend). To measure mTORC1 activity, cells were fixed with 4% paraformaldehyde, permeabilized by ice-cold methanol for 30 mins on ice, and then stained with the primary antibody recognizing phosphor-S6 ribosomal protein (Ser235/236) (D57.2.E, CST), followed by incubating with the secondary APC-coupled AffinityPure F(ab’)2 Fragment Donkey anti-Rabbit IgG antibody (711-606-152, Jackson Immunolab).

### RNA-sequencing and analysis

Total cellular RNA was extracted with the Total RNA Kit II (OMEGA). Library construction and sequencing was performed on a HiSeq or Novaseq 2×150 platform by AZENTA Life Sciences (Suzhou). Sequenced reads were mapped to reference murine genome (mm10 assembly) using bowtie2 with default parameters, and Cufflinks was used to estimate the abundance of transcripts (Langmead and Salzberg, 2012; Trapnell et al., 2010). Quantifications of gene expression were performed using RSEM, and the expression levels of genes in each sample were normalized by means of fragments per kilobase of transcript per million mapped reads (FPKM). These RNA-seq data were deposited in the GEO database with the accession number GSE241746 & GSE241747.

After eliminating absent features (zero counts), differential expressed genes (DEGs) between two groups with 2-3 replicates (one mouse per replicate) were compared *via* DEseq2. Genes were considered to be differentially expressed with the cutoff of fold change ≥ 0.414 and *P* value < 0.05. To identify signaling pathways enriched in different samples, we used R package clusterProfiler and software Gene Set Enrichment Analysis (GSEA) of MSigDB gene sets (Hallmark, KEGG, and GO) (Subramanian et al., 2005; Yu et al., 2012). GSEA was performed for each pairwise comparison using gene lists ranked by the Wald statistic.

### PPIX measurement

Levels of PPIX in cells were analyzed by flow cytometry with excitation at 405 nm and emission at 610/20 nm as described previously (Jang et al., 2015; Tsui et al., 2018).

### Statistical Analysis

The GraphPad Prism 8.0 software (GraphPad) was used to generate graphs and perform statistical analysis. Data were expressed as mean ± SEM. An unpaired nonparametric Mann-Whitney test was utilized to compare differences among groups unless otherwise indicated. Values at *P* < 0.05 were considered significant.

## DATA AVAILABILITY

The RNA-Seq data were deposited in the GEO database with the accession number GSE241746 (review token: uxebqggyvjwfdaj) & GSE241747 (review token: krmbyuiwbtwzpwv). The majority of data supporting the findings of this study are presented within the article and its Supplemental Material. Data that are not directly included are available from the corresponding author upon reasonable request.

## FOOTNOTES

## Acknowledgements

This study was financially supported by the National Natural Science Foundation of China (32170914/81970371/31670888 & 81470564); and Priority Academic Program Development of Jiangsu Higher Education Institutions (PAPD).

## Author contributions

JW conceived and coordinated the study; JW, HG, SZ, XYQ, ZWL, CZ and YZ performed the experiments; JW, HG, SZ and XYQ analyzed the data; XQD conducted bioinformatic analyses and generated relevant figures; HMW, FYG and XMZ assisted in flow cytometry analysis/sorting; JW and YZ drafted the manuscript; JW, XMG and YZ revised the manuscript and secured financial support.

## Conflict of Interest

The authors declare no conflict of interest.

## Materials & Correspondence

material request should be addressed to Dr Jun Wang

---

**Table S2.**
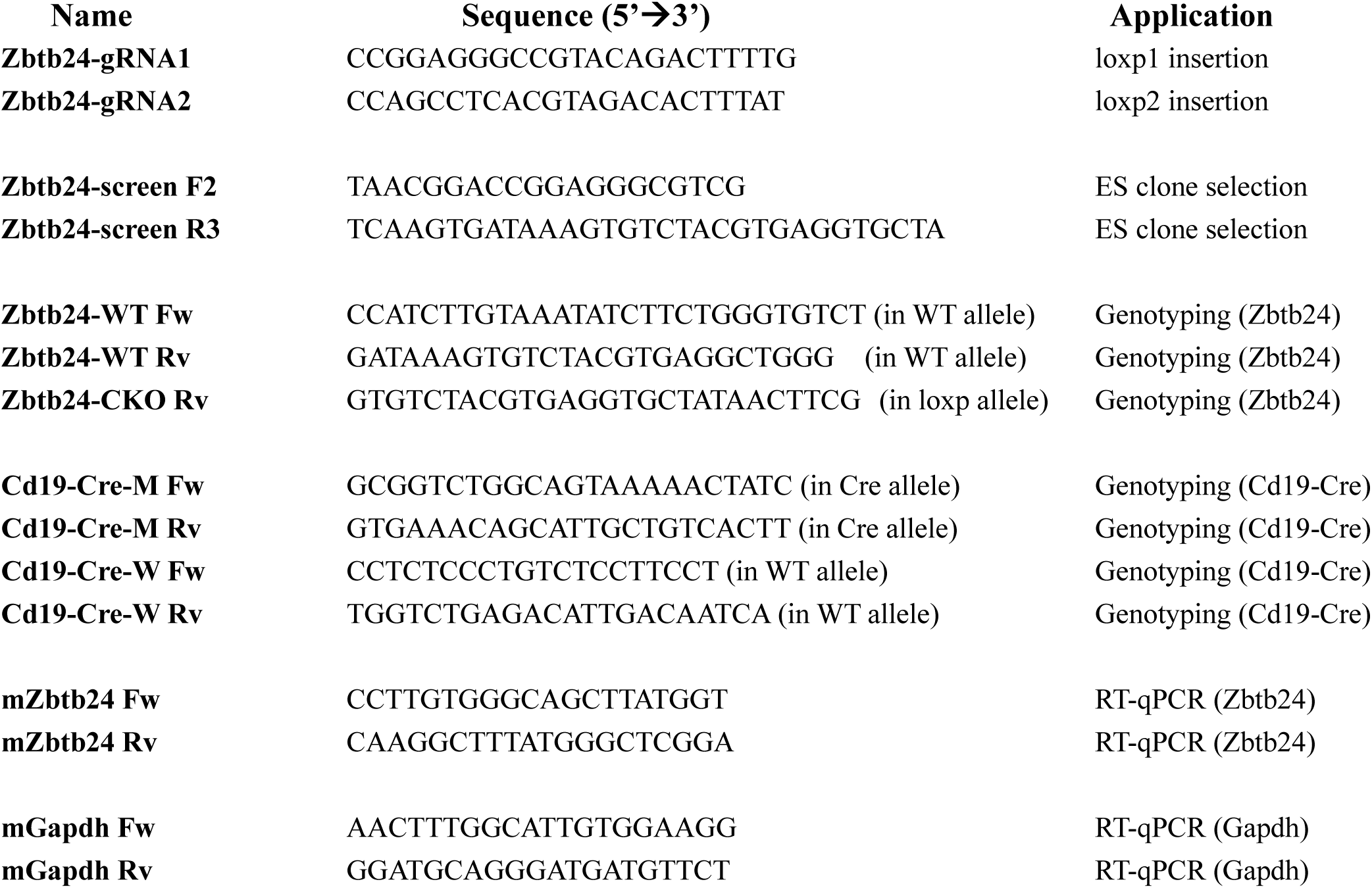
gRNA and primer sequences used in this study.

**Figure S1.**
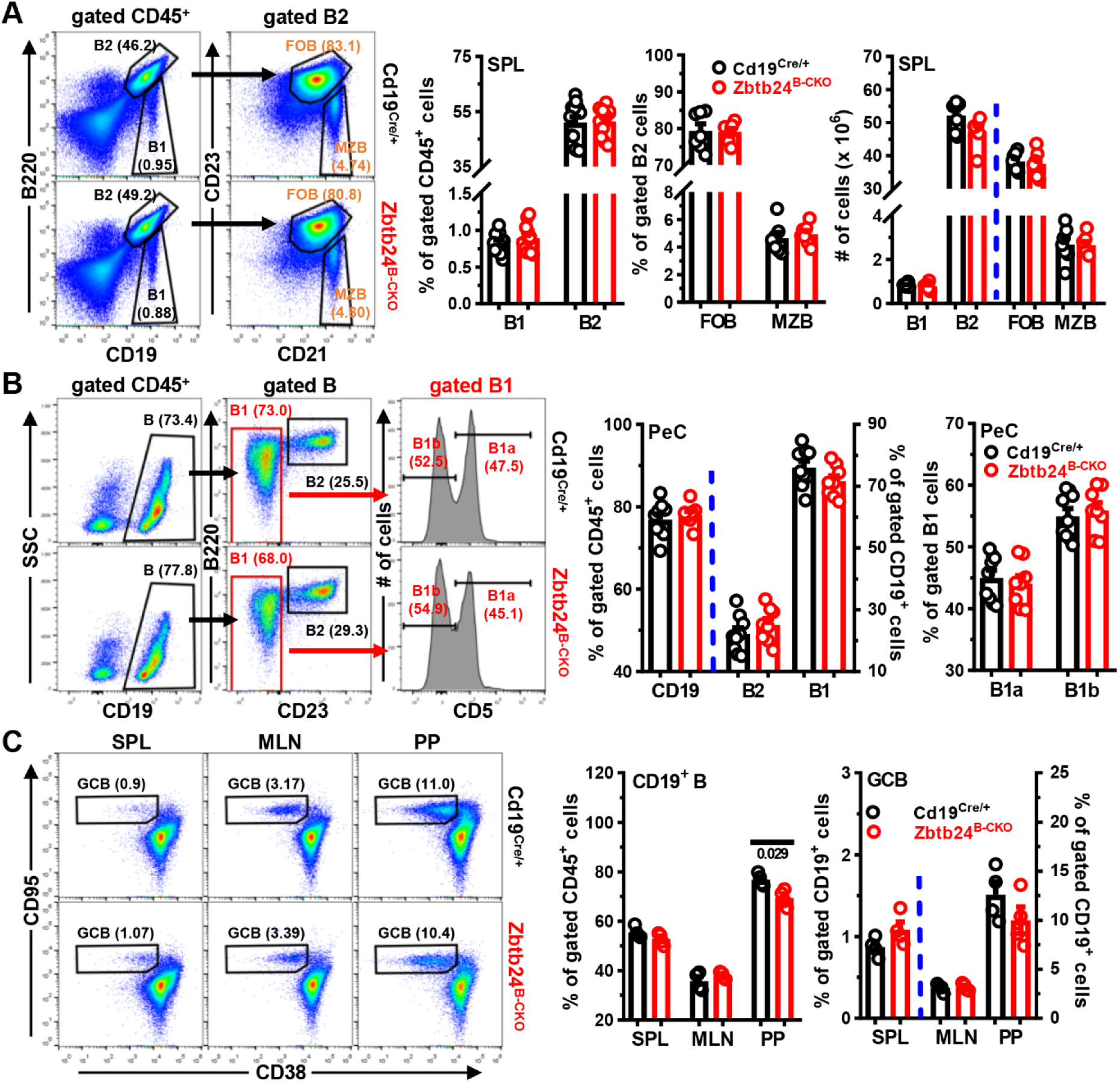
Little impact of Zbtb24-deficiency on the phenotype of B cells in the periphery of mice. Cells from the spleens (SPL), peritoneal cavities (PeC), mesenteric lymph nodes (MLN) and Peyer’s patches (PP) of mice (female, 8-10 weeks old) were stained with antibodies against the indicated surface molecules before flow cytometry analyses. **A&B**, representative pictures/bar graphs showing the gating strategies/percentages of B2 (CD19^+^B220^high^ in SPL or CD19^+^B220^high^CD23^+^ in PeC, respectively), B1 (CD19^+^B220^low^ in SPL or CD19^+^B220^low^CD23^-^ in PeC, respectively), B1a/b (CD5^+^/CD5^-^, respectively, within gated B1), follicular (FOB)/marginzal zone (MZB) (CD23^high^CD21^low^/CD23^low^CD21^high^, repectively, within gated B2) in the SPL (**A**) or PeC (**B**) of Cd19^Cre/+^ and Zbtb24^B-CKO^ mice. The absolute cell numbers are also shown in the far-right panel of **A**. **C**, representative pseudo-plots/bar graphs showing the gating strategies/percentages of germinal center B (GCB, CD95^high^CD38^low^) within gated CD19^+^ B cells in the SPL, MLN and PP of Cd19^Cre/+^ & Zbtb24^B-CKO^ mice. Each symbol represents a single mouse of the indicated genotype. The absolute numbers of indicated B-cell subsets did not differ significantly in the PeC, MLN and PP of the two groups of mice (data not shown). AU, arbitrary units. Pooled data from two-independent experiments were shown in **A&B**.

**Figure S2.**
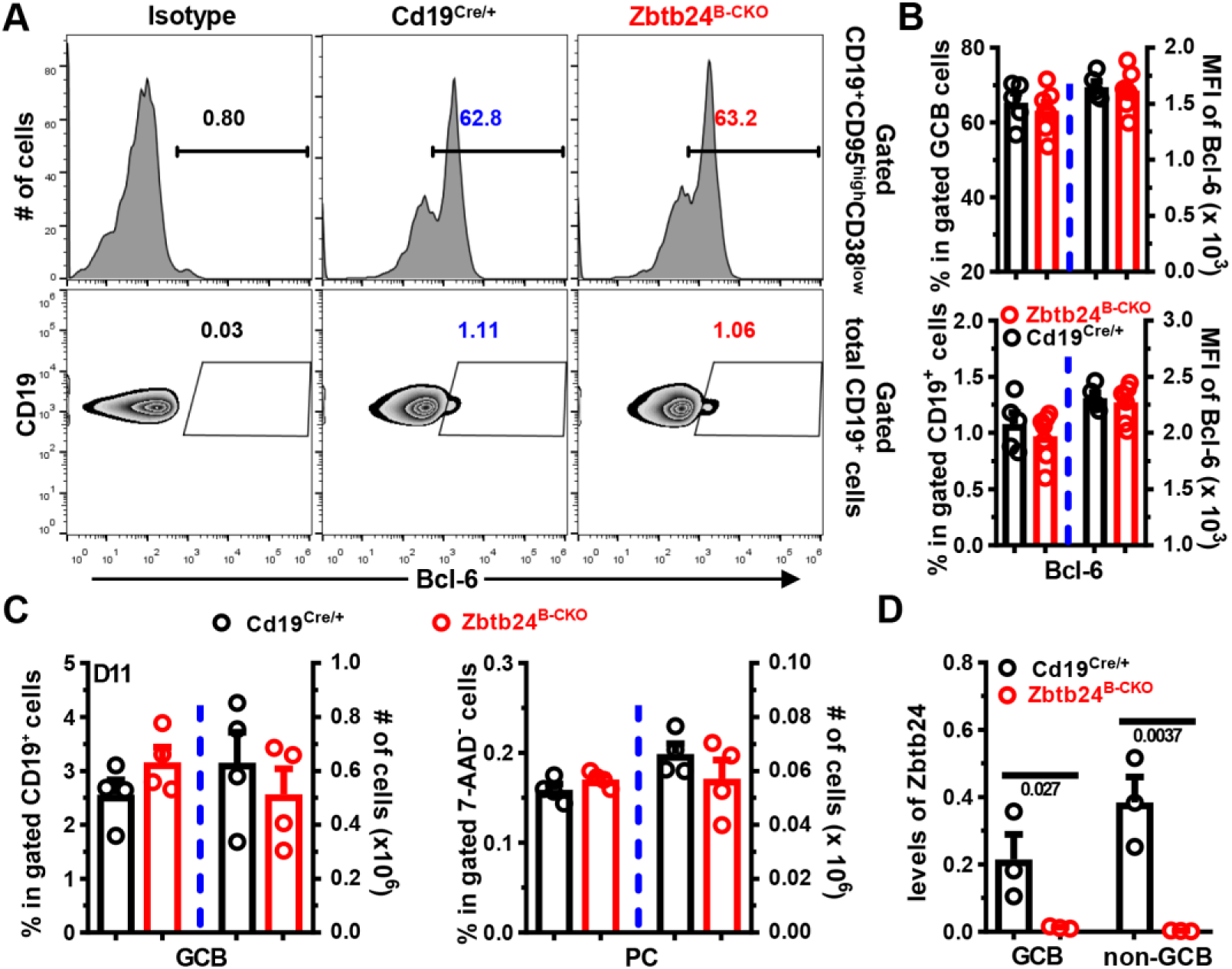
No effect of Zbtb24-deficiency on the phenotype and number of GCB cells after TD-Ag immunization. Splenic cells in mice were stained with antibodies against CD19/CD95/CD38 & Bcl-6 (intracellular) or CD19/CD138 plus 7-AAD to visualize the percents/phenotypes of GCB or PC cells in mice depicted in Figure 2A-C on D14 post NP_19_-OVA/IFA immunization (**A&B**) or on D11 post i.p. immunization with sheep red blood cells (SRBC, 1×10^9^ cells/200 μl PBS/mouse, female, 8 weeks of age) (**C&D**). **A**, representative histograms (*upper panel*) or zebra-plots (*lower panel*) showing the percentages of Bcl6^+^ cells in gated CD19^+^CD95^high^CD38^low^ GCB cells (*upper panel*) or total CD19^+^ B cells (*lower panel*) in control Cd19^Cre/+^ and Zbtb24^B-CKO^ mice. **B**, bar graphs showing the percentages of Bcl6^+^ cells/MFI of Bcl6 in gated CD19^+^CD95^high^CD38^low^ GCB cells (*upper panel*) or total CD19^+^ B cells (*lower panel*). **C**, bar graphs showing the percentages/absolute numbers of CD19^+^CD95^high^CD38^low^ GCB and CD19^low^CD138^high^ PC cells in spleens of mice on D11 post immunization. **D**, mRNA levels of Zbtb24 in FACS-purified CD19^+^CD95^high^CD38^low^ GCB and CD19^+^CD95^-^CD38^+^ non-GCB cells from spleens of immunized mice. Each symbol represents a single mouse of the indicated genotype, and numbers below horizon lines indicate *P* values determined by student *t*-test.

**Figure S3.**
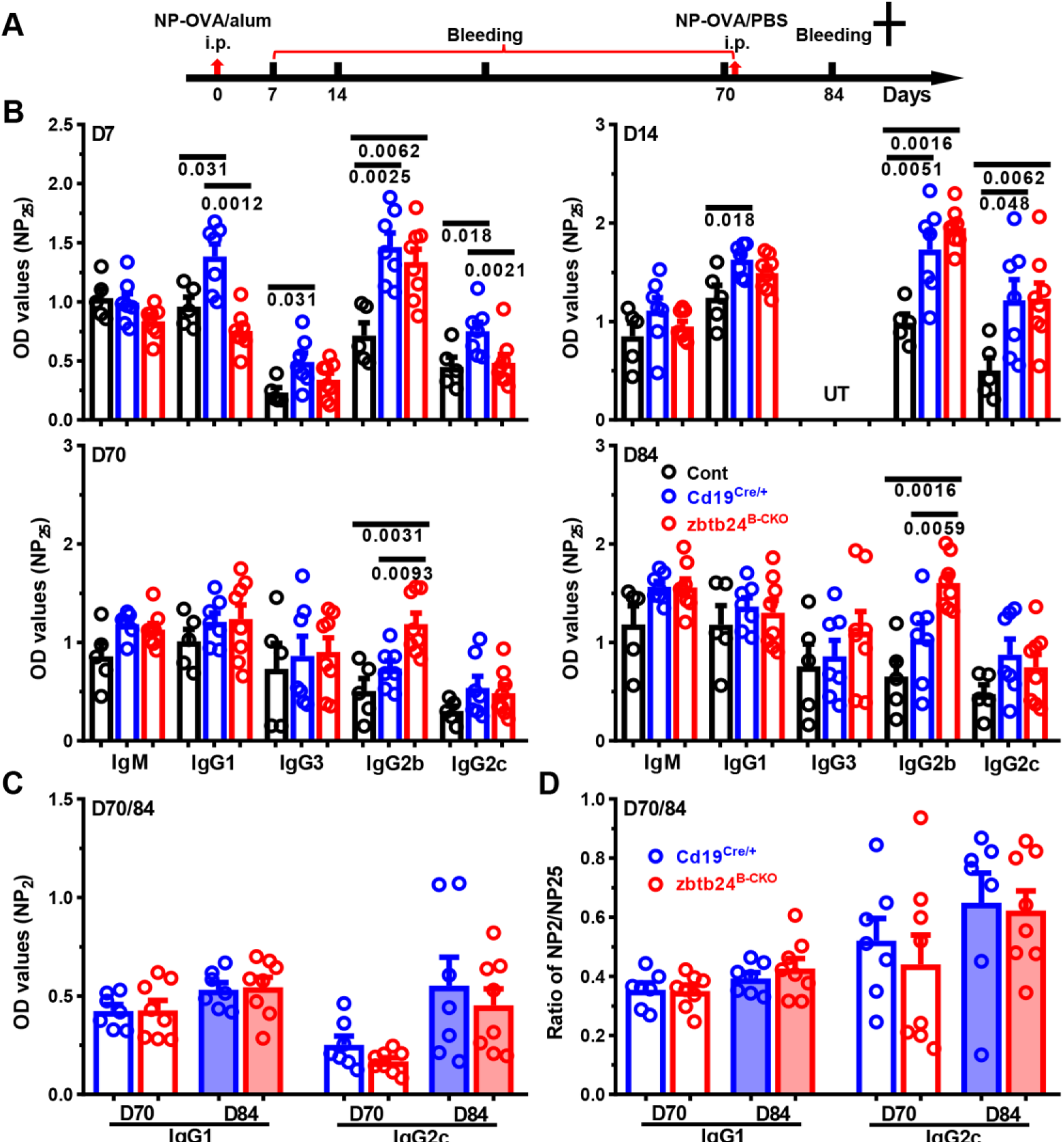
Deficiency of Zbtb24 in B cells has no impact on the long-term antibody responses in mice after NP_19_-OVA/alum immunization. Mice (female, 8 weeks of age) were i.p. immunized with alum-precipitated NP_19_-OVA (NP-OVA/alum, 100 μg/100 μl/mouse) on D0, and rechallenged with NP_19_-OVA in PBS (50 μg/100 μl/mouse) on D70. **A**, a schematic diagram depicting the experimental setup. **B&C**, bar graphs showing the optical density (OD) values of NP-specific antibody subtypes against coated NP_25_-BSA (NP_25_, **B**) or NP_2_-BSA (NP_2_, **C**) in diluted sera of Zbtb24^loxp/loxp^ (Cont), Cd19^Cre/+^ and Zbtb24^B-CKO^ mice on indicated days. **D**, bar graphs showing the ratios of relatively high-affinity (NP_2_) to low-affinity (NP_25_) NP-specific IgG1 & IgG2c in Cd19^Cre/+^ and Zbtb24^B-CKO^ mice on D70 & D84. Each dot represents a single mouse of the indicated genotype and numbers below horizonal lines indicate *P* values determined by Mann-Whitney test. UT, untested.

**Figure S4.**
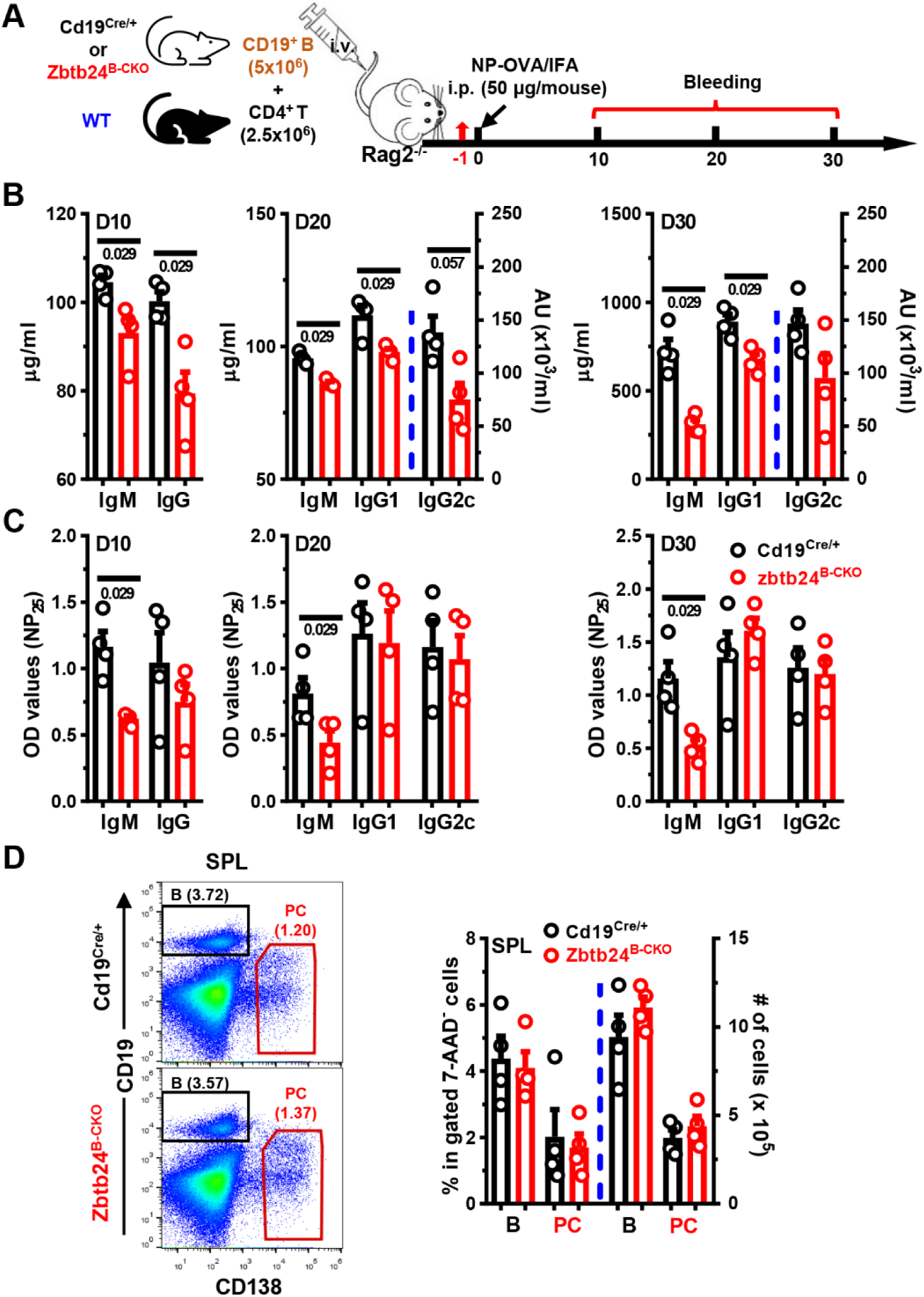
Reduced total antibody-producing ability of Zbtb24^B-CKO^ splenic B cells after adoptive transfer. CD19^+^ B cells, enriched from Zbtb24^B-CKO^ or control Cd19^Cre/+^ mice (male, 8 weeks of age) by magnetic beads, were mixed with CD4^+^ T cells (purified from male WT mice) at the ratio of 2:1 before being intravenously (i.v.) injected into the Rag2^-/-^ recipient mice (5×10^6^ B cells plus 2.5×10^6^ T cells per mouse). One day later, recipient mice were immunized with NP_19_-OVA emulsified in IFA (NP-OVA/IFA) intraperitoneally (i.p.). Blood was taken at indicated times, and total or NP-specific antibody levels in sera were determined by ELISA. **A**, a schematic flow-chart of the experiment setup. **B&C**, bar graphs showing levels of total (**B)** or NP-specific (**C)** IgM/IgG or IgG1/IgG2c levels in sera of recipient mice at indicated times. **E**, bar graphs showing the percentages and absolute numbers of CD19^+^ B cells or CD19^-/low^CD138^+^ plasma cells (PC) in the spleens (SPL) and bone marrows (BM) of Cd19^Cre/+^ *vs.* Zbtb24^B-CKO^ mice. Representative pseudo-plots showing the gates for each population were shown in **D**. Each dot represents a single recipient mouse, and numbers below horizontal lines indicate *P* values. AU, arbitrary units.

**Figure S5.**
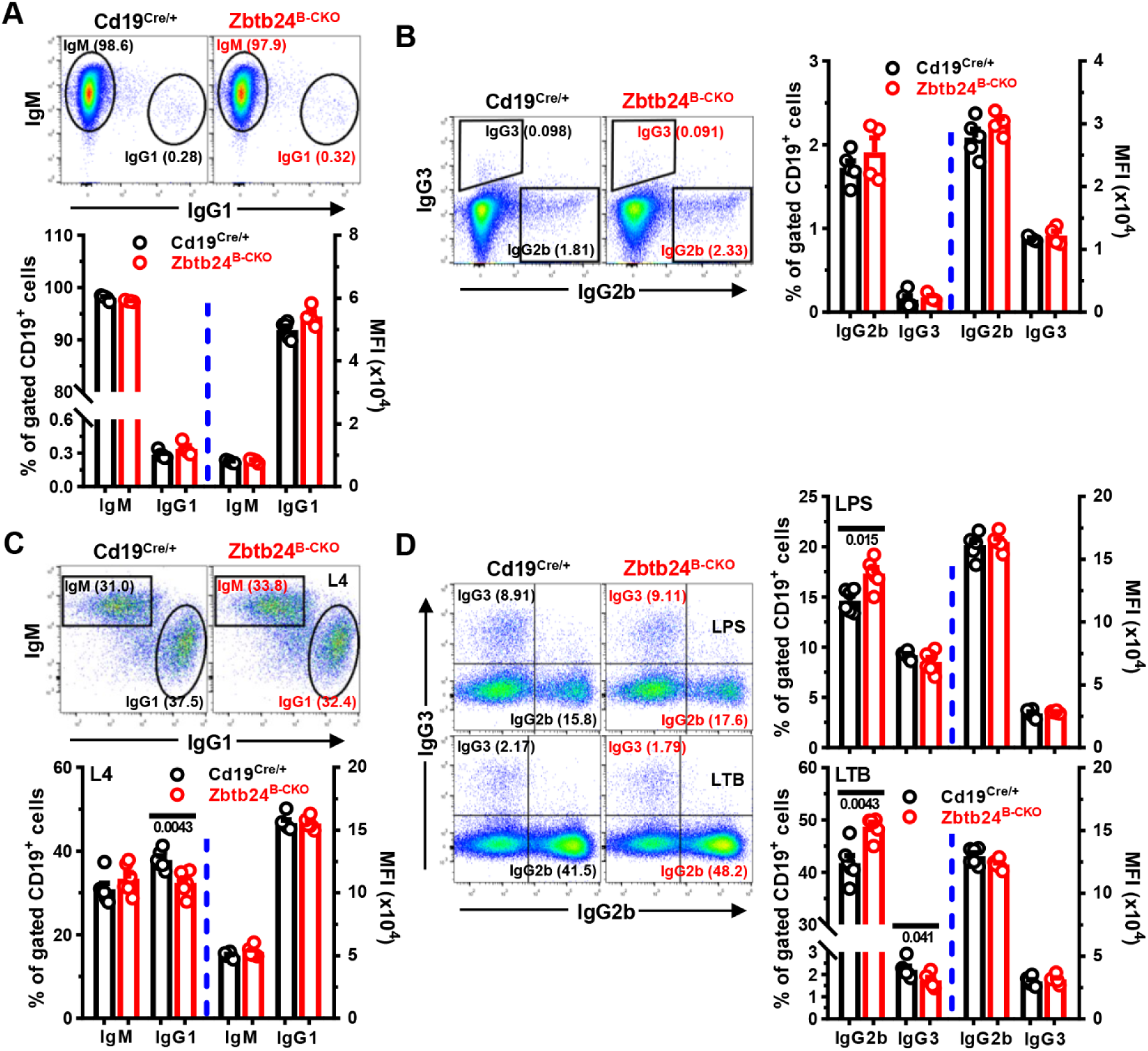
Grossly intact CSR abiltiy of Zbtb24^B-CKO^ B cells. Splenic B cells (2 x 10^5^ cells/well) from Cd19^Cre/+^ or Zbtb24^B-CKO^ mice were stimulated with LPS (10 μg/mL) in the absence/presence of IL-4 (25 ng/ml) or TGF-β (1 ng/ml) plus BAFF (10 ng/ml) in 96 U-bottom plates. On day 4, cells were collected and expressions of surface IgM/IgG2b/IgG3 & IgG1 on B cells were analyzed by flow cytometry. **A&C,** representative pseudo-plots and bar graphs showing the surface IgM/IgG1 on B cells before (**A**) and after culture with LPS plus IL-4 (L4, **C**). **B&D**, representative pseudo-plots/bar graphs showing the surface IgG2b/IgG3 on B cells before (**B**) and after culture with LPS without (*upper panel* in **D**)/with TGF-β & BAFF (LTB, *lower panel* in **D**). Each symbol represents a single mouse of the indicated genotype (female, 10 weeks of age), and numbers below horizontal lines indicate *P* values. MFI, mean fluorescence intensity.

**Figure S6.**
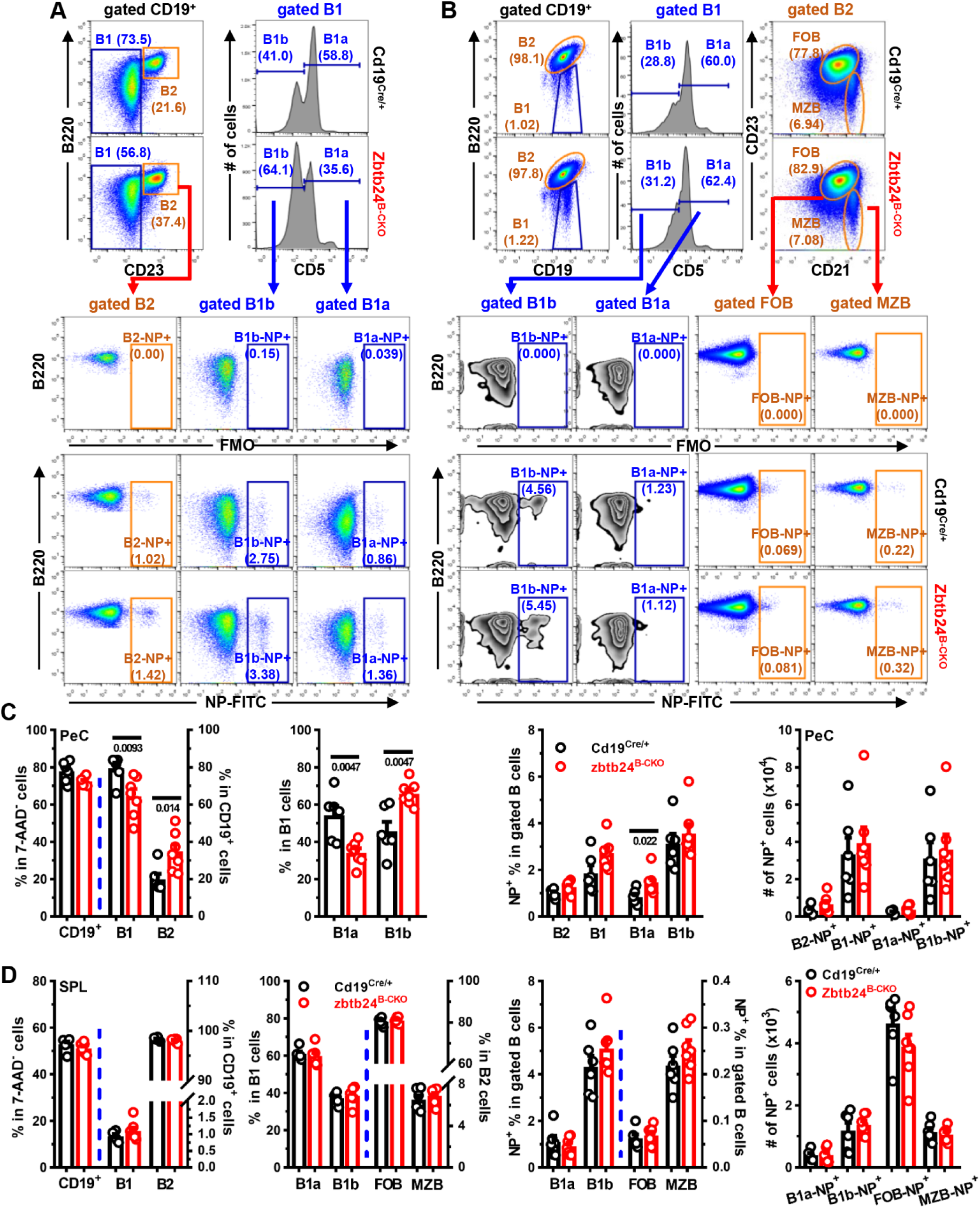
Comparable numbers of NP^+^ B cells in Cd19^Cre/+^ & Zbtb24^B-CKO^ mice on D35 after NP-Ficoll immunization. Peritoneal and splenic cells in NP-Ficoll immunized mice (depicted in Figure 3A-E) were stained with antibodies against B220/CD19/CD5/CD21/CD23 in combination with 7-AAD and NP-FITC. **A&B**, representative flow cytometry plots showing the gating strategies to identify NP^+^ cells within gated B2/B1, B1a/B1b & follicular (FOB)/marginal zone (MZB) B cells in peritoneal cavities (**A**) and spleens (**B**). Gates for NP^+^ cells within B2 (B2-NP+), B1b (B1b-NP+), B1a (B1a-NP+), FOB (FOB-NP+) & MZB (MZB-NP+) were set based on the FMO (fluorescence minus one/FITC-channel) samples. **C&D**, bar graphs showing the percentages/absolute numbers of NP^+^ cells within indicated B-cell subsets in peritoneal cavities (PeC, **C**) and spleens (SPL, **D**) of Cd19^Cre/+^ & Zbtb24^B-CKO^ mice. Each dot represents a single mouse of the indicated genotype, and numbers below horizonal lines indicate *P* values.

**Figure S7.**
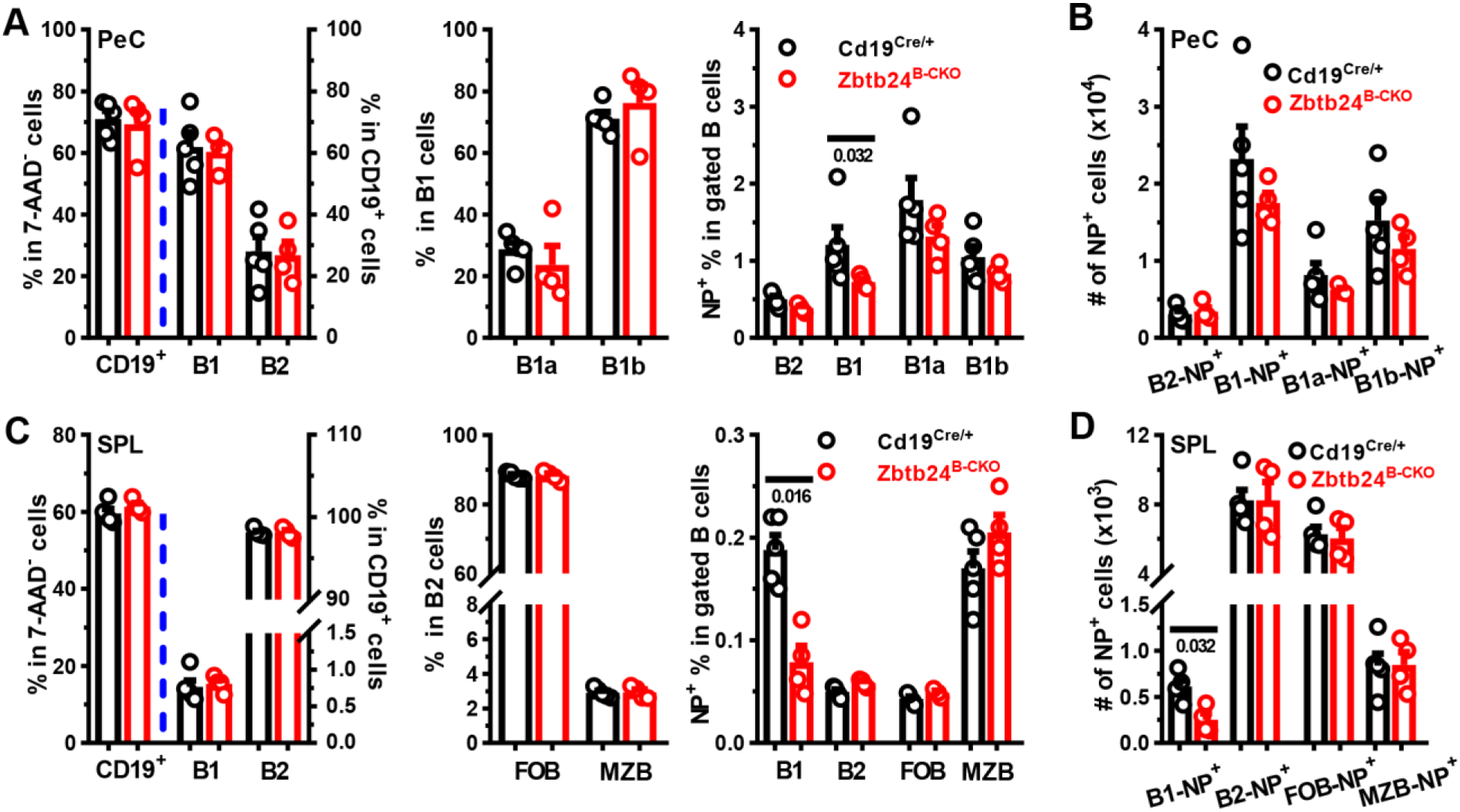
Reduced numbers of NP^+^ B1 cells in Zbtb24^B-CKO^ mice after NP-LPS immunization. Peritoneal and splenic cells in NP-LPS immunized mice (depicted in Figure 3F-H) were stained and analyzed as described in Figure S6A&B. **A&B**, bar graphs showing the percentages (**A**) and absolute numbers (**B**) of NP^+^ cells within indicated B-cell subsets in peritoneal cavities (PeC) of mice. **C&D**, bar graphs showing the percentages (**C**) and absolute numbers (**D**) of NP^+^ cells within indicated B-cell compartments in spleens (SPL) of Cd19^Cre/+^ *vs.* Zbtb24^B-CKO^ mice. Each dot represents a single mouse of the indicated genotype, and numbers below horizonal lines indicate *P* values.

**Figure S8.**
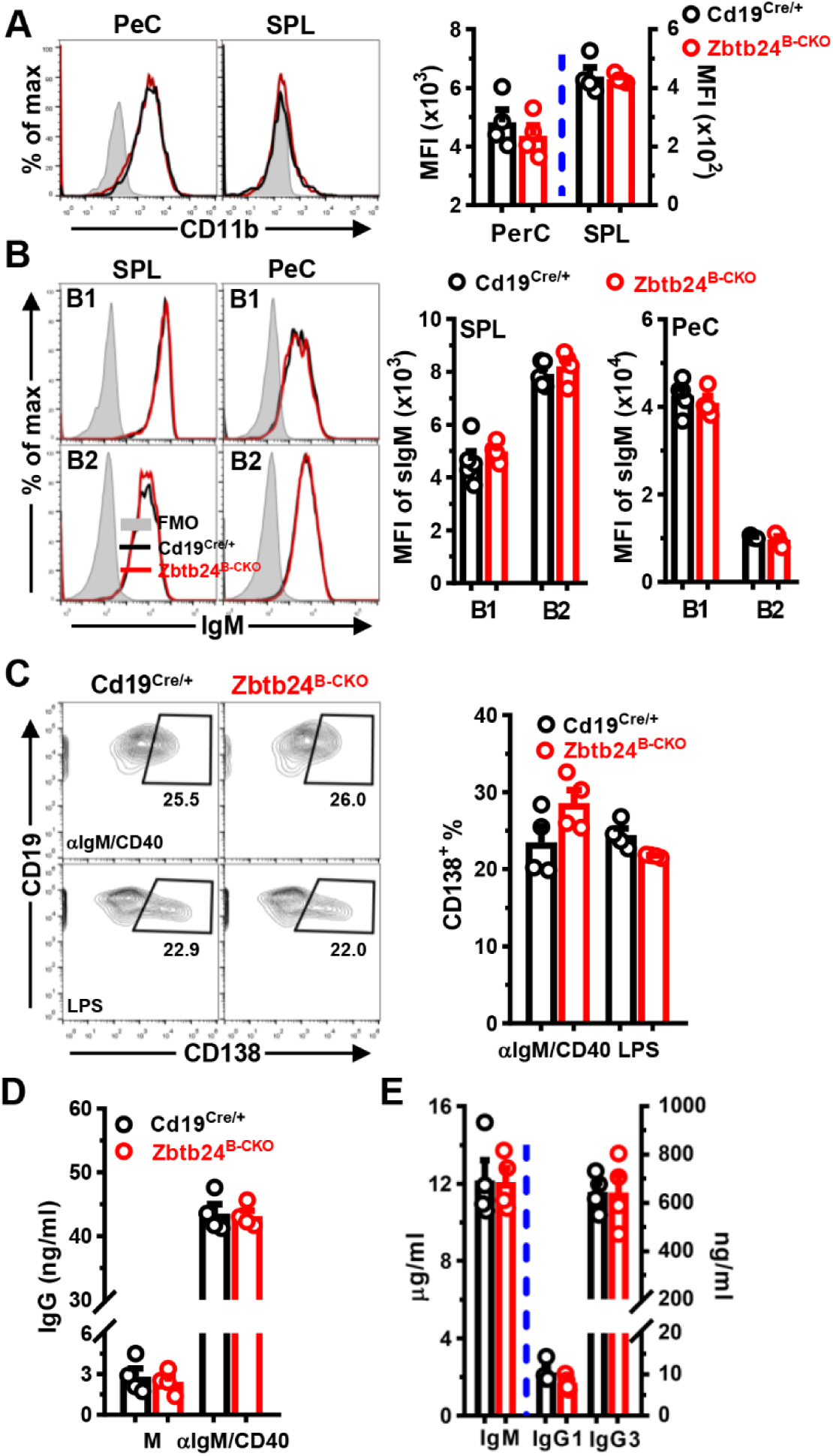
Zbtb24-deficiency has no impact on the differentiation and antibody-producing ability of splenic B cells *in vitro*. **A&B**, representative overlayed-histograms & bar graphs showing levels of surface CD11b (**A**) or IgM (sIgM, **B**) on splenic (SPL)/peritoneal (PeC) B1/B2 cells in Cd19^Cre/+^ *vs.* Zbtb24^B-CKO^ mice. Splenic and peritoneal cells were stained with antibodies against CD19/B220/CD23 & CD11b/IgM before flow cytometry analyses. **C**, representative contour-plots & bar graphs showing percentages of CD138^+^ PC cells in cultured splenic CD19^+^ B cells. **D&E**, levels of IgG (**D**) or IgM/IgG1/IgG3 (**E**) in supernatants of splenic B cells cultured in M, αIgM/CD40 (**D**) or LPS (**E**). CD19^+^ B cells were FACS-sorted from spleens of Cd19^Cre/+^ & Zbtb24^B-CKO^ mice, and subsequently cultured (1 x 10^5^ cells/well in 96-well plate) in medium (M), anti-IgM & anti-CD40 (αIgM/CD40, 5/5 μg/ml) or LPS (10 μg/ml). On day 4, surface expressions of CD138 on cultured B cells were analyzed by flow cytometry, and antibody levels in culture supernatants were determined by ELISA (**C-E**). Each symbol represents a single mouse of the indicated genotype (female, 8-10 weeks of age).

**Figure S9.**
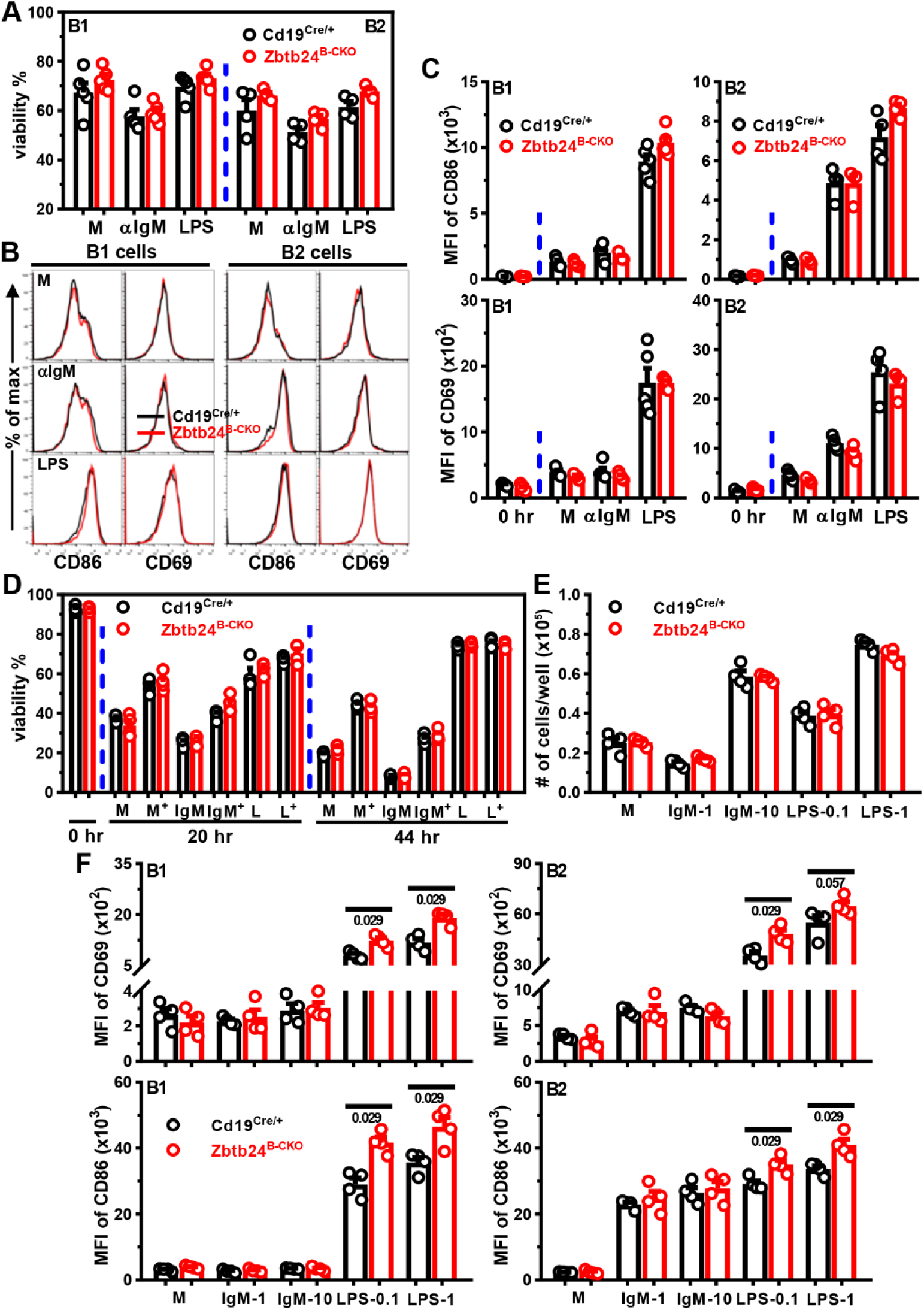
Zbtb24-deficiency does not compromise the survival, activation and proliferation of Zbtb24^B-CKO^ B cells *in vitro*. CD19^+^B220^low^CD23^-^ B1 and CD19^+^B220^high^CD23^+^ B2 cells were FACS-sorted from peritoneal cavities of Cd19^Cre/+^ & Zbtb24^B-CKO^ mice, and subsequently cultured (2-5 x 10^4^ cells/well in 96-U bottom plate) in medium (M), αIgM (2 μg/ml) or LPS (0.5 μg/ml). Viabilities (based on forward/side scatter) and surface levels of CD69/CD86 were analyzed by flow cytometry (**A-C**). **A**, bar graphs showing viabilities of peritoneal B1 & B2 cells at 24 hrs. **B&C**, representative overlayed histograms (**B**) or bar graphs (**C**) showing surface expressions of CD86 & CD69 on 24-hr cultured peritoneal B1/B2 cells derived from Cd19^Cre/+^ *vs.* Zbtb24^B-CKO^ mice before (0 hr) and after 24 hr’s culture. CD19^+^ B cells were purified from spleens of Cd19^Cre/+^ & Zbtb24^B-CKO^ mice, and subsequently cultured (1.5-2 x 10^5^ cells/well in 96-well plate) in medium, anti-IgM or LPS before flow cytometry analysis (**D-F**). **D**, bar graphs showing the percents of living cells (7-AAD^-^ & Annexin V^-^) before (0 hr) or after cultures in M, 1 μg/ml anti-IgM (IgM) or LPS (L) in the absence/presence of 0.2 μg/ml BAFF (indicated by a ‘+’ in the upper-right corner) for 20 & 44 hrs. **E**, bar graphs showing the numbers of living cells (based on forward/side scatter) in different cultures on day 3. **F**, bar graphs showing the expression levels of surface CD69 (top row) and CD86 (bottom row) on gated B2 (CD19^+^B220^high^, left column) or B1 (CD19^+^B220^low^, right column) cells after 20 hrs’ culture in medium (M), 1/10 μg/ml anti-IgM (IgM-1/IgM-10, respectively), 0.1/1 μg/ml LPS (LPS-0.1/LPS-1, respectively). Each symbol represents a single mouse of the indicated genotype (12-week-old males in **A-C**, and 8-week-old females in **D-F**), and numbers below horizontal lines in **F** indicate *P* values.

**Figure S10.**
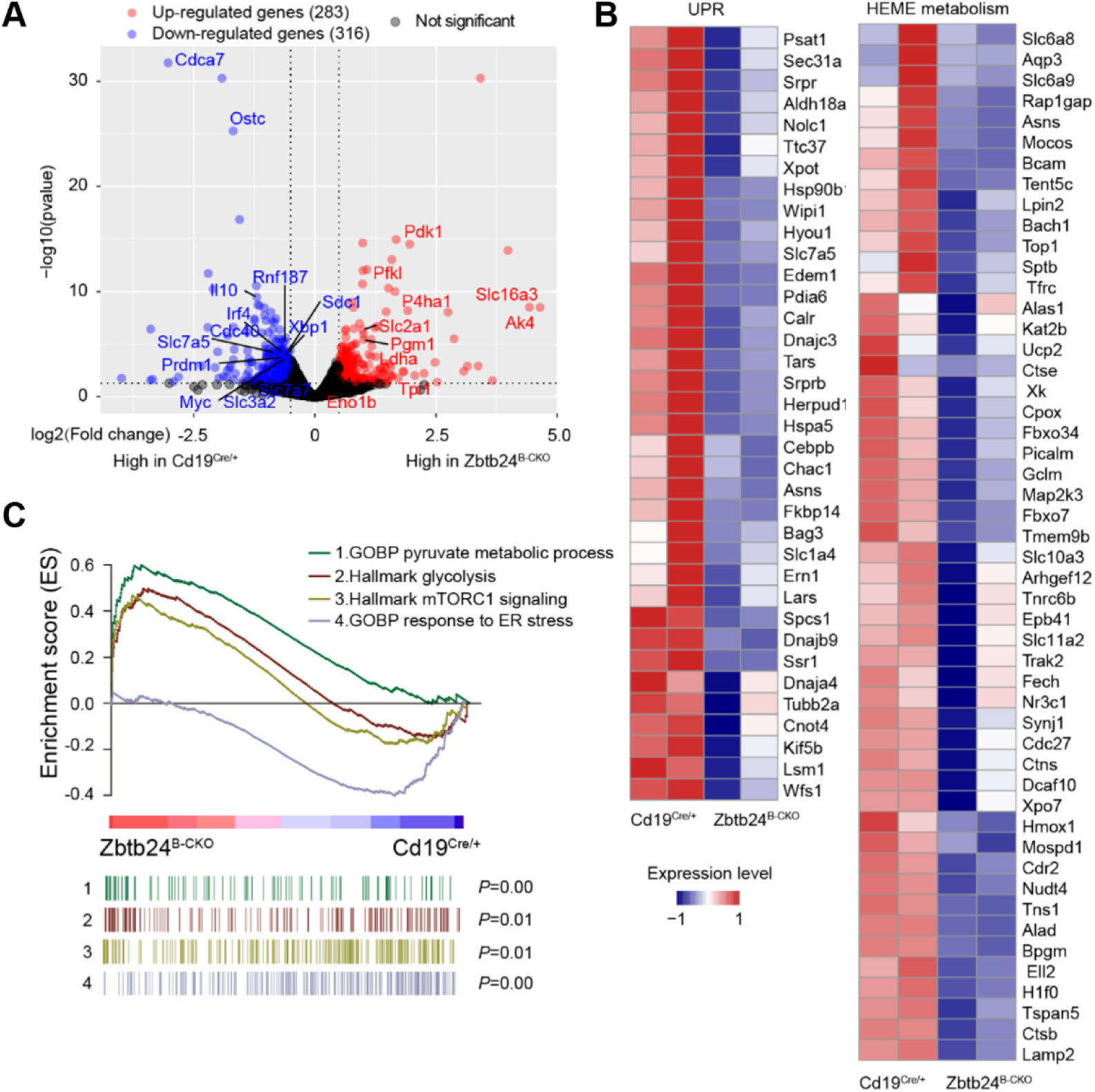
Dysregulated genes/signaling pathways in Zbtb24-deficient B1 cells. FACS-sorted peritoneal CD19^+^B220^low^CD23^-^ B1 cells were stimulated with 0.1 μg/ml LPS in 96-U bottom plate for 24 hrs (related to Figure 6A-C). **A**, volcano plots showing upregulated (red) or downregulated (blue) genes in Zbtb24^B-CKO^ B1 cells. **B**, heatmaps showing the enriched genes affiliated to UPR and heme metabolism in Zbtb24-deficient B1 cells by GSEA. **C**, GSEA plots showing the enrichment of genes regulating pyruvate metabolism, glycolysis, mTORC1 signaling and ER stress in LPS-stimulated Zbtb24^B-CKO^ *vs.* Cd19^Cre/+^ B1 cells.

**Figure S11.**
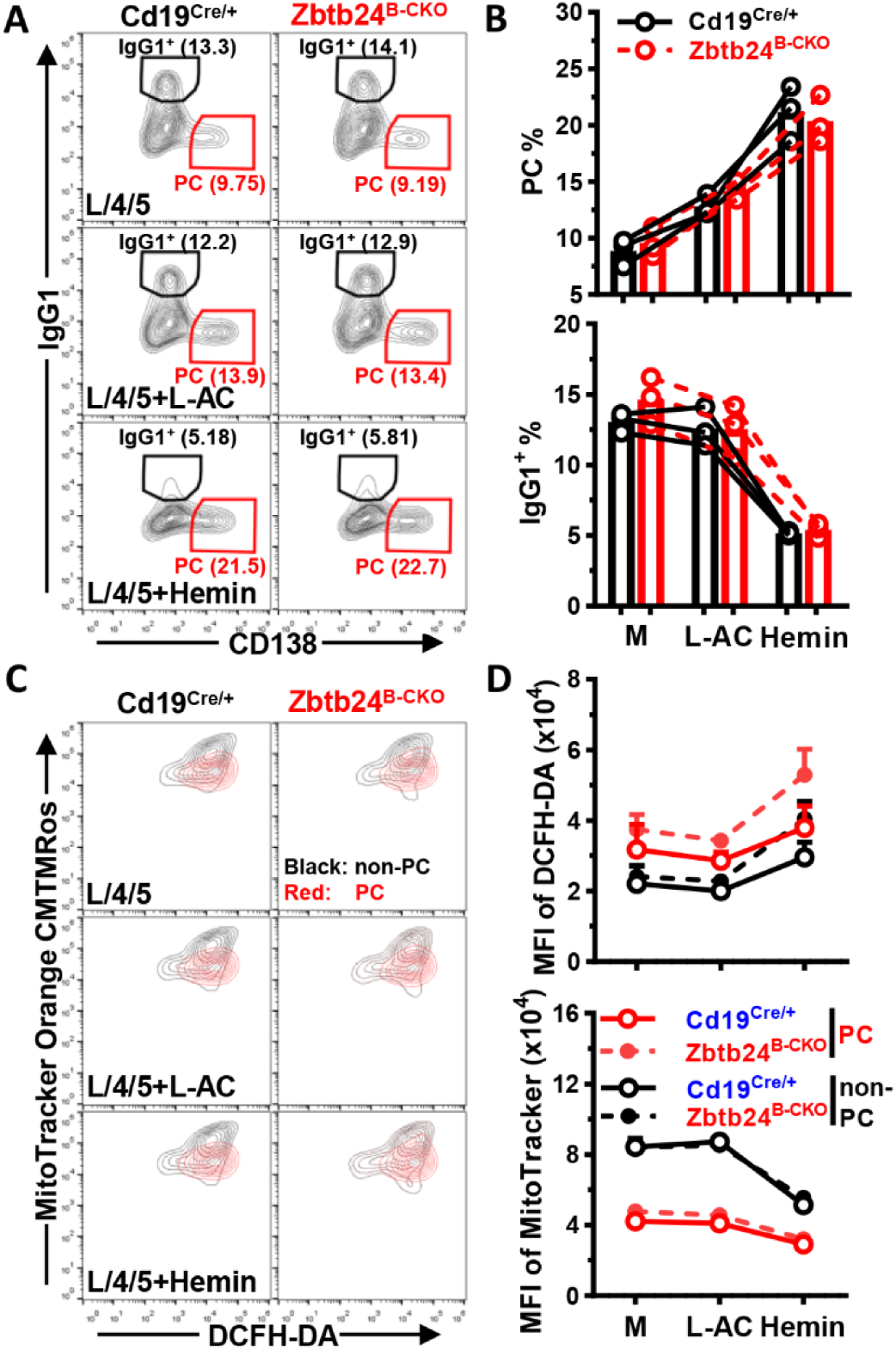
No effect of Zbtb24-deficiency on heme metabolism in stimulated splenic B cells. Splenic B cells, isolated from Cd19^Cre/+^ or Zbtb24^B-CKO^ mice, were stimulated with 10 μg/ml LPS plus 10 ng/ml IL-4 & IL-5 (L/4/5) without/with additional L-AC (250 μM) or Hemin (25 μM) in 96-U bottom plate for 4 days. **A&B**, representative contour-plots (**A**) and bar graphs (**B**) showing the percentages of class-switched CD138^-^IgG1^+^ B cells (IgG1^+^) or differentiated IgG1^-^CD138^+^ PC in stimulated Cd19^Cre/+^ *vs.* Zbtb24^B-CKO^ splenic B cells on day 4. **C**, representative overlayed contour-plots showing the intracellular ROS levels (visualized by DCFH-DA) and mitochondrial mass/membrane potentials (detected by MitoTracker Orange CMTMRos) in gated CD138^-^ non-PC (black) *vs.* CD138^+^ PC (red) in cultured splenic B cells on day 4. **D**, line graphs showing the ROS levels (MFI of DCFH-DA, *upper panel*) and mitochondrial mass/membrane potential (MFI of MitoTracker, *lower panel*) in gated non-PC/PC cells in Cd19^Cre/+^ *vs.* Zbtb24^B-CKO^ splenic B cell cultures. Each dot in **B** represents a single mouse of the indicated genotype (male, 14 weeks of age), while results in **D** are expressed as mean ± SEM (n=3).

## Notes

### Competing Interest Statement

The authors have declared no competing interest.

### Summary of Updates

The Distribution/Reuse Options were changed.

